# Crosstalk between eIF2α and eEF2 phosphorylation pathways optimizes translational arrest in response to oxidative stress

**DOI:** 10.1101/694752

**Authors:** Marisa Sanchez, Yingying Lin, Chih-Cheng Yang, Philip McQuary, Alexandre Rosa Campos, Pedro Aza Blanc, Dieter A. Wolf

## Abstract

The cellular stress response triggers a cascade of events leading to transcriptional reprogramming and a transient inhibition of global protein synthesis, which is thought to be mediated by phosphorylation of eukaryotic initiation factor-2α (eIF2α). Using mouse embryonic fibroblasts (MEFs) and the fission yeast S. pombe, we report here that rapid translational arrest and cell survival in response to hydrogen peroxide-induced oxidative stress do not rely on eIF2α kinases and eIF2α phosphorylation. Rather H_2_O_2_ induces a block in elongation through phosphorylation of eukaryotic elongation factor 2 (eEF2). Kinetic and dose-response analyses uncovered crosstalk between the eIF2α and eEF2 phosphorylation pathways, indicating that, in MEFs, eEF2 phosphorylation initiates the acute shutdown in translation, which is then maintained by eIF2α phosphorylation. Our results challenge the common conception that eIF2α phosphorylation is the primary trigger of translational arrest in response to oxidative stress and point to integrated control that may facilitate the survival of cancer cells.

**HIGHLIGHTS:**

- Oxidative stress-induced translation arrest is independent of eIF2α phosphorylation
- Oxidative stress blocks translation elongation
- Oxidative stress triggers eEF2 kinase activation
- eEF2K KO cells are hypersensitive to oxidative stress

## INTRODUCTION

mRNA translation, the energetically most demanding step in gene expression, is tightly regulated (Buttgereit and Brand, 1995; Rolfe and Brown, 1997; Sonenberg and Hinnebusch, 2009). Although translation is conceptually separated into discrete biochemical steps - initiation, elongation, termination, and ribosome recycling (Hershey et al., 2012) – regulation may occur in a more integrated fashion in vivo, affecting multiple steps simultaneously (Richter and Coller, 2015; Sha et al., 2009).

One of the best-studied regulatory pathways centers around eukaryotic initiation factor 2 (eIF2), a GTP-binding protein containing three subunits, α, β and γ. As part of the ternary complex (also comprising of the initiator tRNA-methionine and GTP), eIF2 joins the 40S ribosome in scanning the mRNA for an initiation codon. After the 60S ribosomal subunit is added upon start codon recognition, GTP is hydrolyzed, leading to the release of eIF2-GDP. To be reused in subsequent rounds of initiation, eIF2 bound to GDP must be converted to eIF2-GTP by the guanine nucleotide exchange factor eIF2B. Phosphorylation of eIF2α on residue S51 increases its affinity for eIF2B but reduces its GDP to GTP exchange activity thus resulting in reduced levels of functional eIF2 leading to inhibition of global mRNA translation (Pavitt, 2018).

eIF2α phosphorylation is pivotal for ablating translation in response to environmental stress (Wek et al., 2006). Eukaryotic cells recognize and process diverse stress signals to elicit programs of gene expression that are designed to alleviate cellular damage, or alternatively induce apoptosis. Important contributors to this stress response are a family of four protein kinases (PERK, PKR, GCN2, and HRI) that phosphorylate eIF2α on Ser51 and inhibit protein synthesis, thereby conserving energy and facilitating the reprogramming of gene expression and signaling to restore protein homeostasis (Wek, 2018; Wek et al., 2006). However, previous reports in yeast showed that the eIF2α kinase Gcn2 as well as the eIF2α phosphorylation site are dispensable for translational inhibition in response to H_2_O_2_ and ultraviolet light (Knutsen et al., 2015; Shenton et al., 2006). These studies also provided evidence that H_2_O_2_ impedes translation elongation, although the underlying signaling and the interplay between effects on initiation and elongation remained unclear. Likewise, it remained open whether control at the level of elongation is conserved in mammalian cells.

Even though initiation of translation is considered the most important regulatory step, there is increasing attention on mechanisms that affect other steps, in particular elongation (Richter and Coller, 2015). Two important elongation regulatory factors are eukaryotic elongation factor 2 (eEF2) and its kinase, eEF2K. Eukaryotic elongation factor 2 kinase (eEF2K) is a Ca^2+^/calmodulin (CaM)-dependent kinase, which negatively modulates protein synthesis by phosphorylating eEF2 (Kenney et al., 2014; Tavares et al., 2014). Phosphorylation at Thr56 within the GTP-binding domain of eEF2 prevents its recruitment to ribosomes and thus blocks elongation (Carlberg et al., 1990). eEF2K is subject to regulation by cellular nutrient and energy status. Nutrient depletion is associated with activation of eEF2K via AMPK and inhibition of TORC1 signaling pathways, resulting in increased autophagy and cell survival (Kruiswijk et al., 2012). The frequent overexpression of eEF2K in human cancers may thus confer tumor cell adaptation to microenvironmental nutrient stress (Fu et al., 2014). As inhibition of eEF2K’s survival function augments ER stress-induced apoptosis, eEF2K may be an anti-cancer drug target (Cheng et al., 2013; Fu et al., 2014).

The tumorigenic state is marked by alterations activating a mutually reinforcing network of metabolic, genotoxic, proteotoxic, and oxidative stresses (Gorrini et al., 2013; Luo et al., 2009). Oxidative stress is particularly significant because it can both induce and result from the other forms of stress. Moderate levels of reactive oxygen species (ROS) may contribute to tumorigenesis either as signaling molecules or by inducing DNA mutations (Gorrini et al., 2013). To sustain survival under high ROS levels typically seen in tumors, cells rely on sophisticated stress defense pathways that have been proposed as cancer drug targets (Luo et al., 2009). In particular, overwhelming oxidative stress defense either through inhibition of ROS scavenging or through augmenting ROS levels by chemotherapeutics or inducers of protein misfolding has been proposed as a promising strategy (Gorrini et al., 2013; Trachootham et al., 2009; Wolf, 2014).

The important roles of ROS-mediated oxidative stress in cancer promotion and therapy contrast with the limited understanding of the effects of ROS on global gene expression programs that direct cell adaptation or death decisions. Whereas effects on transcription, mediated through prominent transcription factors such as NFκB, AP1, and NRF2, are well documented (Trachootham et al., 2008), post-transcriptional levels of control remain largely unassessed. The prevailing dogma that stress impacts gene expression post-transcriptionally through shutdown of global translation as a consequence of eIF2α phosphorylation (Wek, 2018; Wek et al., 2006) has been qualified by studies in yeast (Knutsen et al., 2015; Shenton et al., 2006). To gain further insight into post-transcriptional control of gene expression in response to oxidative stress, we have addressed mechanisms conserved in mammalian cells and in fission yeast. Our data suggest that an integrated response at the levels of translation initiation and elongation coordinates survival under oxidative stress.

## RESULTS

### Oxidative stress-induced transcriptome changes differ depending on the status of eIF2α phosphorylation

To gain insight into layers of gene expression in response to oxidative stress, we obtained transcriptomic profiles by RNA sequencing. Immortalized mouse embryonic fibroblasts (MEFs) were used that carried either wildtype eIF2α (WT) or a point mutant in which the inhibitory Ser 51 phosphorylation site was replaced by alanine (eIF2α^S51A^) thus rendering eIF2α insensitive to kinase-mediated inhibition and translational shutdown in response to stress (Back et al., 2009; McEwen et al., 2005). To monitor acute and prolonged responses to oxidative stress, cells were exposed for either 15 or 120 minutes to 500 μM H_2_O_2_, and triplicate RNA samples were obtained for sequencing. Although eIF2α^S51A^ MEFs had ~2-fold higher levels of cellular reactive oxygen species (ROS) at baseline, treatment with 500 μM H_2_O_2_ led to a robust and comparable increase in ROS levels in both cell lines (Figure S1A,B).

Significant H_2_O_2_-induced changes in mRNA levels (p ≤ 0.05) at both treatment time points were identified with 613 mRNAs changing in wildtype MEFs and 982 in eIF2α^S51A^ mutant MEFs. There was partial overlap in the responsive mRNAs but an approximately equal number of changes were unique to each cell line (Figure S1C). In WT cells, the transcriptional response peaked at 15 min (Figure 1A) and mostly consisted of downregulated mRNAs that encoded factors involved in protein synthesis (Figure 1B). Many of these changes were already present in eIF2α^S51A^ MEFs at baseline (untreated, 0 min and Figure S1D,E), indicating that the mutant cells display a partially activated stress response, possibly as a result of higher basal ROS levels (Figure S1A,B). Since mRNAs encoding protein synthesis factors, including ribosomal proteins, are typically among the most highly translated in unstressed cells, their acute downregulation may serve to liberate ribosome capacity for the translation of newly induced mRNAs (Lackner et al., 2012; Lee et al., 2011). After 120 min, WT cells induced mRNAs encoding factors involved in protein folding, signifying the recovery phase (Figure 1B).

**Figure 1:**
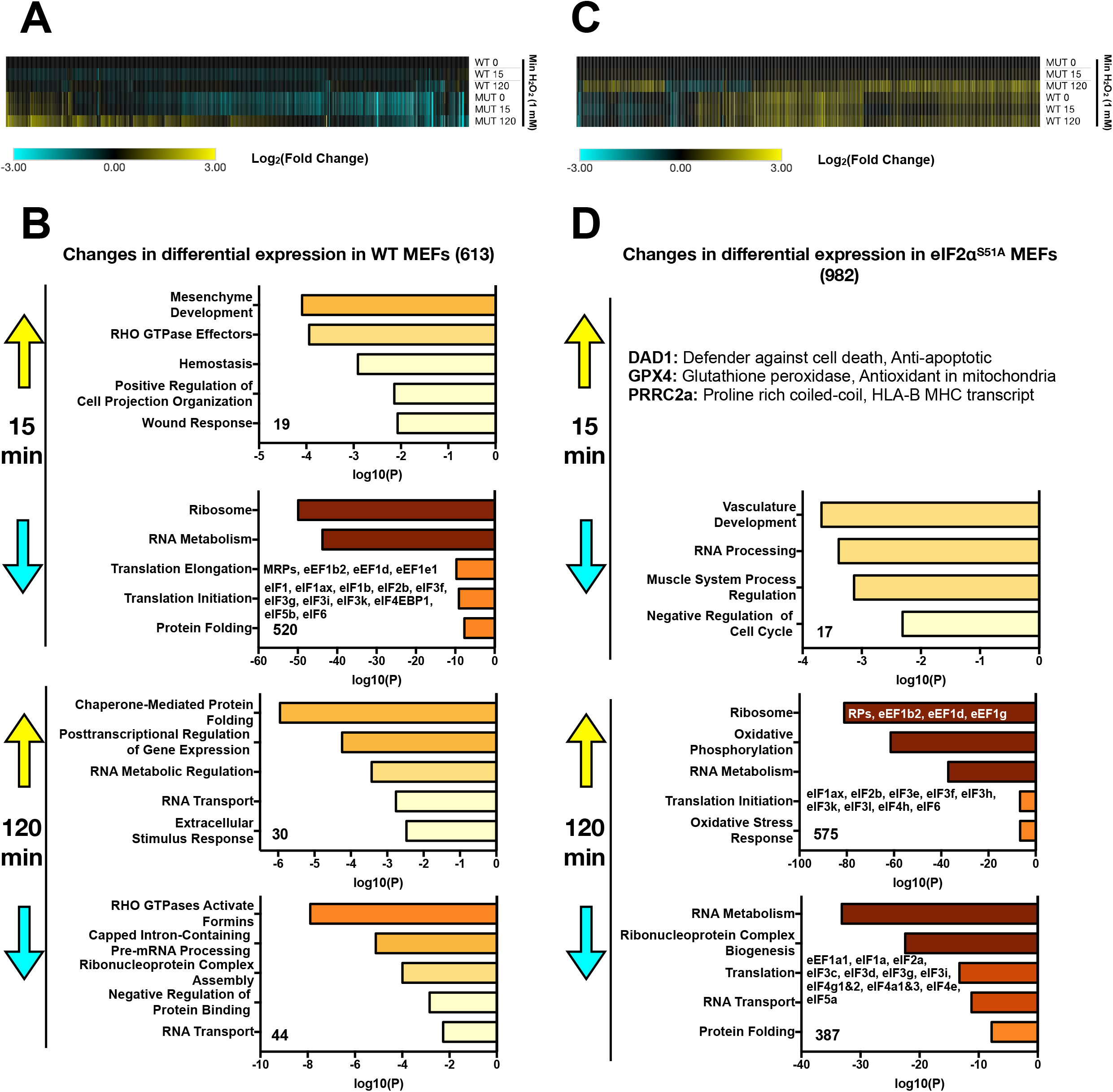
Transcriptome changes under oxidative stress. (**A**) WT and eIF2α^S51A^ MEFs were either untreated or treated with H_2_O_2_ (500 μM) for 15 min or 120 min. Total RNA was extracted and analyzed by RNAseq. The data represents a compilation of significant changes in expression (p ≤ 0.05) found in WT MEFs at either the 15 or 120 min time point. Results are normalized to untreated WT MEFs at time 0 and displayed as a clustered heat map. Upregulated transcripts are shown in yellow and downregulated transcripts in blue. (**B**) Individual list of H_2_O_2_-induced and H_2_O_2_-repressed mRNAs (±3-fold change, p ≤ 0.05) in WT MEFs were loaded into Metascape and the pathways enriched are indicated at each time point. Colored boxes represent either upregulated (yellow) or downregulated (blue) pathways. (**C**) WT and eIF2α^S51A^ MEFs were either untreated or treated with H_2_O_2_ (500 μM) for 15 min or 120 min. Total RNA was extracted and analyzed by RNAseq. The data represents a compilation of significant changes in expression (p ≤ 0.05) found in eIF2α^S51A^ MEFs at either the 15 or 120 min time point. Results are normalized to untreated eIF2α^S51A^ MEFs at time 0 and displayed as a clustered heat map. Upregulated transcripts are shown in yellow and downregulated transcripts in blue. (**D**) Individual list of H_2_O_2_-induced and H_2_O_2_-repressed mRNAs (±3-fold change, p ≤ 0.05) in eIF2α^S51A^ MEFs were loaded into Metascape and the pathways enriched are indicated at each time point. Colored boxes represent either upregulated (yellow) or downregulated (blue) pathways.

In contrast, few changes were observed in eIF2α^S51A^ mutant MEFs after 15 min and the response was considerably stronger after 120 min (Figure 1C), indicating that eIF2α^S51A^ mutant MEFs show an overall delayed response to H_2_O_2_. Unlike WT cells, which displayed mostly a gene repressive pattern, eIF2α^S51A^ mutant MEFs showed a higher number of mRNAs upregulated than downregulated. Curiously, the pattern in eIF2α^S51A^ MEFs at 120 min was an approximate mirror image of the pattern observed with WT cells at 15 min. Whereas the latter downregulated protein synthesis factors, they were upregulated in eIF2α^S51A^ MEFs (Figure 1D). Conversely, protein folding was up in WT cells but down in eIF2α^S51A^ MEFs. In summary, the mRNA profiles indicate that eIF2α phosphorylation substantially shapes gene expression in stressed and unstressed cells. The observed differences in the regulation of mRNAs encoding translation and protein folding factors may reflect distinct translational responses possibly resulting in differential sensitivity to H_2_O_2_.

### Oxidative stress-induced translational inhibition is independent of eIF2α phosphorylation

To test the above idea, WT and eIF2α^S51A^ mutant MEFs were exposed to inducers of oxidative and ER stress, followed by determination of cell viability. Whereas there was differential sensitivity to the ER stress inducer thapsigargin with eIF2α^S51A^ mutant cells being more sensitive than WT MEFs as described previously (Hiramatsu et al., 2014), both MEF lines showed equal sensitivity to cytotoxicity exerted by hydrogen peroxide (H_2_O_2_) (Figure 2A,B). Likewise, equal sensitivity of both MEF lines was observed for the oxidant tert-butyl hydroperoxide (TBHP).

**Figure 2:**
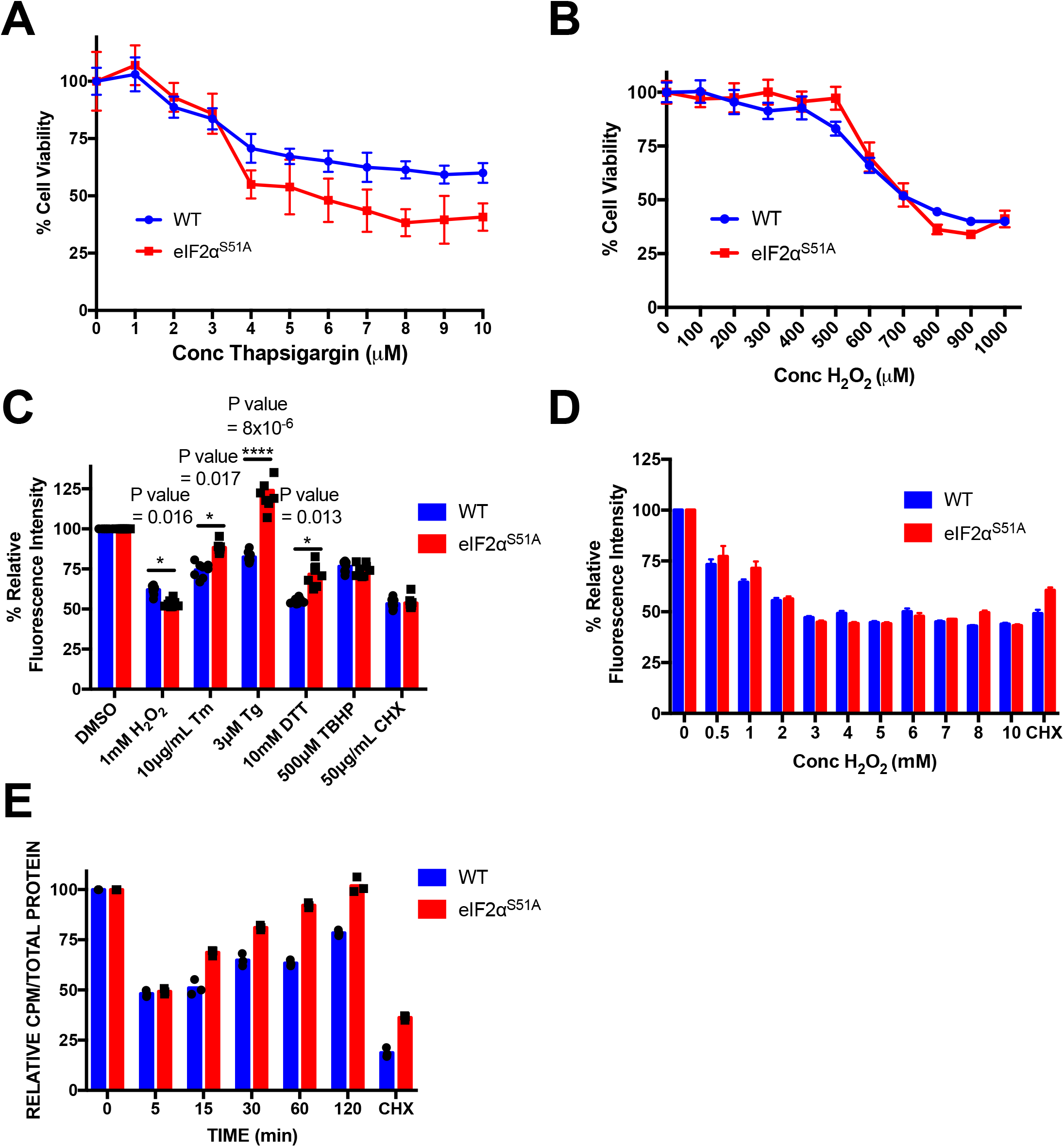
Effect of hydrogen peroxide on viability and protein synthesis. (**A**) Wildtype and eIF2α^S51A^ mutant MEFs were treated with increasing doses (1-10 μM) of the ER stress inducer thapsigargin for 2 hours. MTT assay was used to assess cell viability. The graph represents the mean ± standard deviations of 8 replicates. (**B**) Wildtype and eIF2α^S51A^ mutant MEFs were treated with the indicated increasing doses (100-1000 μM) of the oxidative stress inducer hydrogen peroxide (H_2_O_2_) for 2 hours. MTT assay was used to assess cell viability. The graph represents the mean ± standard deviations of 8 replicates. (**C**) Wildtype and eIF2α^S51A^ mutant MEFs were treated with the indicated doses of specific stress inducers: H_2_O_2_, tunicamycin (Tm), thapsigargin (Tg), dithiothreitol (DTT) and tert-butyl hydroperoxide (TBHP). Cells were treated for 2 hours except in the case of tunicamycin, which was added for 4 hours. The protein synthesis inhibitor cycloheximide (CHX) was used as a control. Protein synthesis was measured by fluorescently tagging nascent polypeptides and quantifying the fluorescence intensity. The graph represents the normalized mean ± standard deviations of eight replicates. The individual symbols represent individual data points. *P value H_2_O_2_ = 0.016, P value Tm = 0.017, P value Tg = 8 x 10^−6^, P value DTT = 0.013. (**D**) Wildtype and eIF2α^S51A^ mutant MEFs were treated with increasing doses (0.5-10 mM) of H_2_O_2_ for 1 hour. Protein synthesis was measured with a fluorescent tagging assay. 50 μg/mL CHX was added for 1 hour as a control. The graph represents the normalized mean ± standard deviations of eight replicates. (**E**) Wildtype and eIF2α^S51A^ mutant MEFs were treated with 500 μM H_2_O_2_ for increasing periods (5 - 120 min). Cells were simultaneously labeled with [^35^S]-methionine for 5 minutes. Protein synthesis was measured by quantifying radioactive counts per minute (CPM) and adjusted according to the total amount of protein. The graph represents the normalized mean ± standard deviations of at least two independent experiments each done in triplicates. The individual symbols represent individual data points.

The same differential response was observed at the level of global protein synthesis. Whereas wildtype MEFs downregulated incorporation of fluorescently labeled amino acids into cellular proteins in response to ER stress inducers tunicamycin (Tm), thapsigargin (Tg), and dithiothreitol (DTT), eIF2α^S51A^ mutant MEFs were less responsive (Figure 2C). In contrast, both wildtype and eIF2α^S51A^ mutant MEFs responded with a ~50% inhibition of amino acid incorporation to challenge with 1 mM hydrogen peroxide (Figure 2C). An equal ~50% inhibition was obtained for both cell lines with the ribosomal elongation blocker cycloheximide (Figure 2C).

Likewise, dose responses for inhibition of protein synthesis by hydrogen peroxide did not differ between wildtype and eIF2α^S51A^ mutant MEFs with both cell lines showing maximal inhibition at a concentration of 3 mM H_2_O_2_ (Figure 2D). However, time course measurements with brief 5 minute pulses of [^35^S]-methionine revealed markedly different kinetics of recovery of protein synthesis from insult with 1 mM H_2_O_2_. Whereas both cell lines exhibited acute inhibition of protein synthesis within 5 minutes of exposure to hydrogen peroxide, eIF2α^S51A^ mutant MEFs recovered protein synthesis sooner than wildtype MEFs, returning to pretreatment levels within 60 minutes (Figure 2E). Recovering wildtype MEFs did not reach pretreatment levels within the 120-minute time frame of the experiment.

While this data confirmed the canonical role of eIF2α phosphorylation as a mediator of ER stress-induced translation inhibition, it revealed an unexpected differential response to oxidative stress. Whereas eIF2α phosphorylation is dispensable for immediate translational shutdown following H_2_O_2_, it is required for maintenance of the inhibition, potentially enabling time demanding repair and recovery processes.

### eIF2α kinases are dispensable for translational shutdown and survival under oxidative stress

In order to gain a more mechanistic understanding of eIF2α phosphorylation-independent translation inhibition under oxidative stress, we investigated the role of eIF2α kinases. For this, we turned to the fission yeast S. pombe as a model system due to its facile genetics and faithful recapitulation of the complexity of eIF2α phosphorylation. Like human cells, S. pombe encodes several different eIF2α kinases (Hri1p, Hri2p, Gcn2p) that phosphorylate a conserved serine residue (S52). Strains either carrying a S52A mutation of eIF2α or deleted for all three eIF2α kinases were fully viable as described previously (Berlanga et al., 2010; Tvegård et al., 2007). Remarkably, both strains showed similar inhibition of protein synthesis in response to H_2_O_2_ as observed in wildtype cells despite being completely deficient in eIF2α phosphorylation (Figure 3A,B).

**Figure 3:**
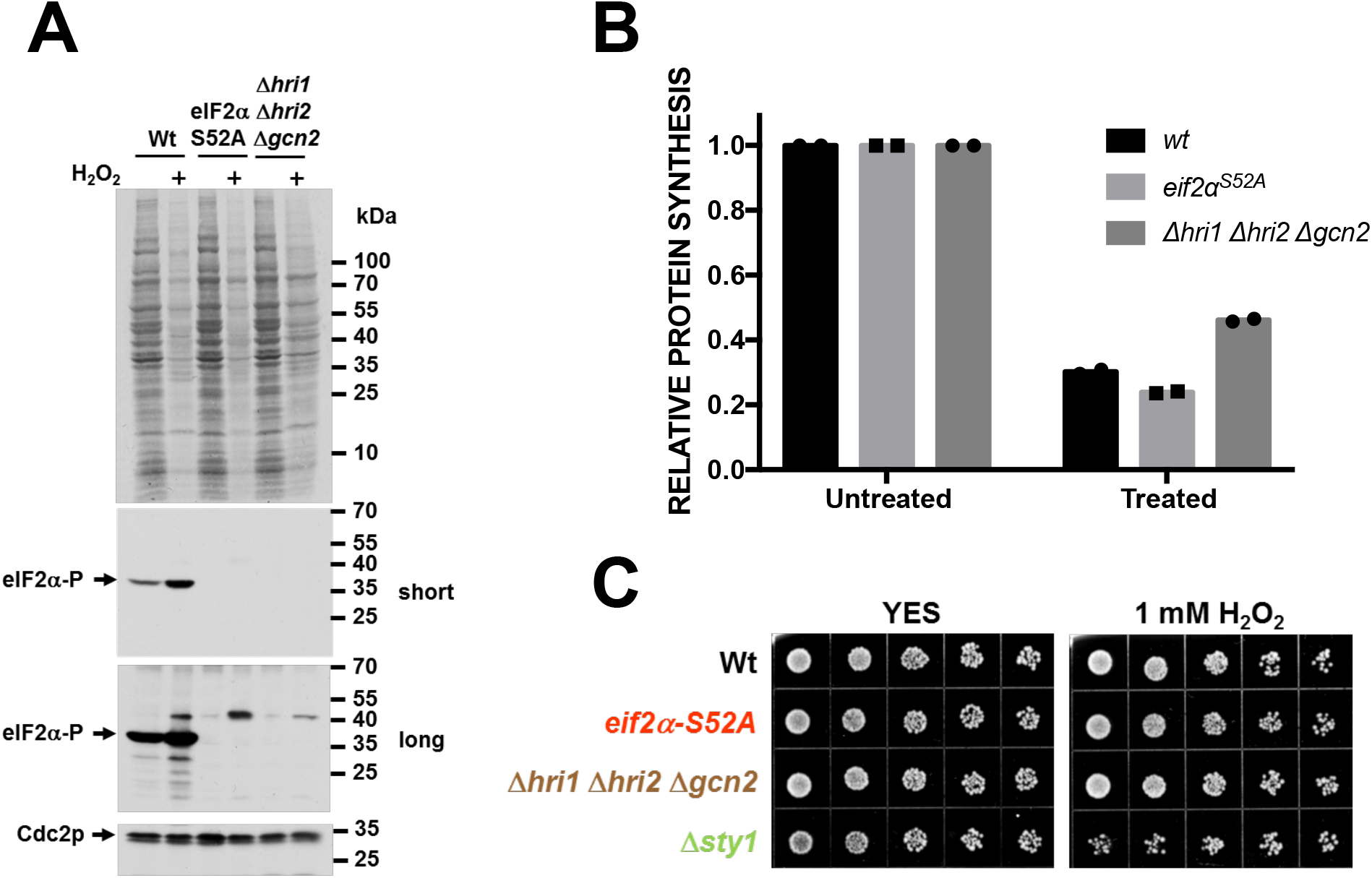
Effect of S. pombe eIF2α phosphorylation pathway mutations on protein synthesis and cell survival. (**A**) WT, *eIF2α^S52A^*, and *Δhri1 Δhri2 Δgcn2* strains of *S. pombe* were treated with 1 mM H_2_O_2_ for 15 minutes and [^35^S]-methionine was added for the next 45 minutes. Total protein was isolated, separated by SDS-PAGE and protein synthesis was detected by film autoradiography (top panel). The same lysates were analyzed by immunoblotting for eIF2α phosphorylation. Signals for Cdc2p are shown for reference. (**B**) The same lysates as described in (A) were TCA precipitated and [^35^S]-methionine incorporation was quantified by scintillation counting. (**C**) Sensitivity of WT, *eIF2α^S52A^, Δhri1 Δhri2 Δgcn2* and *Δsty1* strains to H_2_O_2_ was assessed by spotting 5-fold serial dilutions on plates containing 1 mM H_2_O_2_.

Consistent with the data on inhibition of protein synthesis, neither the eIF2α S52A mutant nor the triple eIF2α kinase deficient strain was sensitive to H_2_O_2_ exposure as compared to a strain deleted for the stress responsive MAP kinase, Sty1p (Figure 3C, S3). In summary, these data strongly suggest that the conserved eIF2α phosphorylation site as well as the kinases modifying it are dispensable for both acute translational shutdown in response to oxidative stress and cell survival.

### Translation elongation is blocked in the absence of eIF2α phosphorylation under oxidative stress

To corroborate these findings, we determined the effect of H_2_O_2_ on the global distribution of ribosomes along a sucrose density gradient. In wildtype cells, H_2_O_2_ triggered a time-dependent redistribution of ribosomes from the polysomal to the monosomal 80S fraction, indicating that cells responded to oxidative stress with polysome run-off and suppression of the formation of new polysomes (Figure 4A). Only a marginal polysome to monosome shift was observed in eIF2α^S51A^ mutant MEFs under the same conditions (Figure 4A). This finding suggested that ribosomes do not run off mRNAs in eIF2α^S51A^ mutant MEFs exposed to oxidative stress.

**Figure 4:**
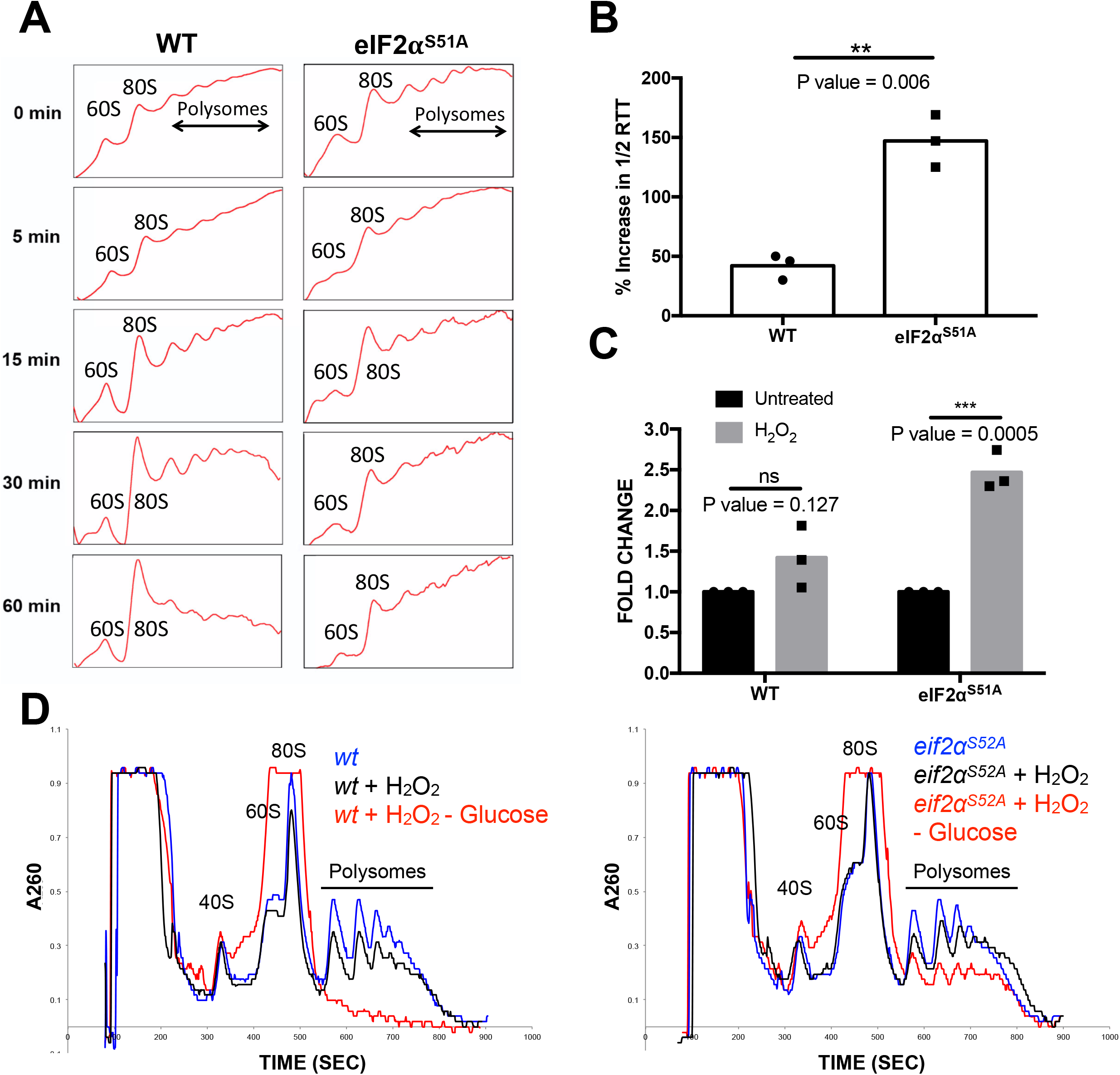
The effects of oxidative stress on translation elongation. (**A**) Polysome profiling was done with wildtype and eIF2α^S51A^ mutant MEFs treated with 500 μM H_2_O_2_ for increasing periods (5 – 60 min). (**B**) Ribosome transit times were analyzed in wildtype and eIF2α^S51A^ mutant MEFs treated with 500 μM H_2_O_2_. The graph represents the mean percent increase in 1/2 transit times ± standard deviations of at least three independent experiments each done in triplicates. *P value = 0.006. (**C**) The graph represents the normalized mean fold change of 1/2 transit times ± standard deviations of the same experiments as shown in Figure 3B. *P value WT = 0.127, P value eIF2α^S51A^ = 0.0005. (**D**) Polysome profiles of wildtype and eIF2α^S52A^ mutant *S. pombe.* Cells were either untreated or treated with 1 mM H_2_O_2_ for 1 hour followed by removal of glucose for 5 minutes.

Persistent polyribosome occupancy on mRNA between 5 and 60 minutes after exposure of eIF2α^S51A^ mutant MEFs to H_2_O_2_ was in stark contrast to the marked inhibition of amino acid incorporation after only 5 minutes of H_2_O_2_ (Figure 2E). This seemingly conflicting data led to the hypothesis that oxidative stress induced a block of translation elongation. Indeed, ribosome transit time (Nielsen and McConkey, 1980) was ~2.5-fold increased in H_2_O_2_-treated eIF2α^S51A^ mutant MEFs relative to wildtype MEFs, which showed less than a 1.5-fold increase (Figure 4B,C).

To further substantiate an H_2_O_2_-induced translation elongation block, wildtype and eIF2α^S52A^ mutant strains of *S. pombe* were subjected to brief glucose withdrawal to induce polysome run-off ((Ashe et al., 2000); Figure S4C). After 1 hour of exposure to H_2_O_2_, cells showed a decrease in polysomes that was more pronounced in wildtype cells relative to eIF2α^S52A^ mutant cells (Figure 4D). Glucose depletion of H_2_O_2_ treated wildtype cells led to complete polysome run-off (Figure 4D). In contrast, eIF2α^S52A^ cells maintained substantial polysome levels under the same conditions, indicating reduced translation elongation (Figure 4D). Taken together, this data strongly suggested that oxidative stress induces a block in translation elongation that is especially apparent in cells unable to phosphorylate eIF2α.

### eEF2K signaling is upregulated under H_2_O_2_-induced oxidative stress in the absence of eIF2α phosphorylation

We next assessed the potential involvement of negative regulators of translation elongation, specifically the kinase that phosphorylates translation elongation factor eEF2 (eEF2K). Wildtype and eIF2α^S51A^ mutant MEFs were subjected to various ER and oxidative stress inducers followed by assessment of the inhibitory phosphorylation of eEF2 on threonine 56. eEF2 phosphorylation was induced in both wildtype and eIF2α^S51A^ mutant MEFs treated with H_2_O_2_, TBHP, and DTT (Figure 5A). However, eEF2 phosphorylation in response to oxidants, particularly H_2_O_2_, was considerably more pronounced in eIF2α^S51A^ mutant MEFs (Figure 5A).

**Figure 5:**
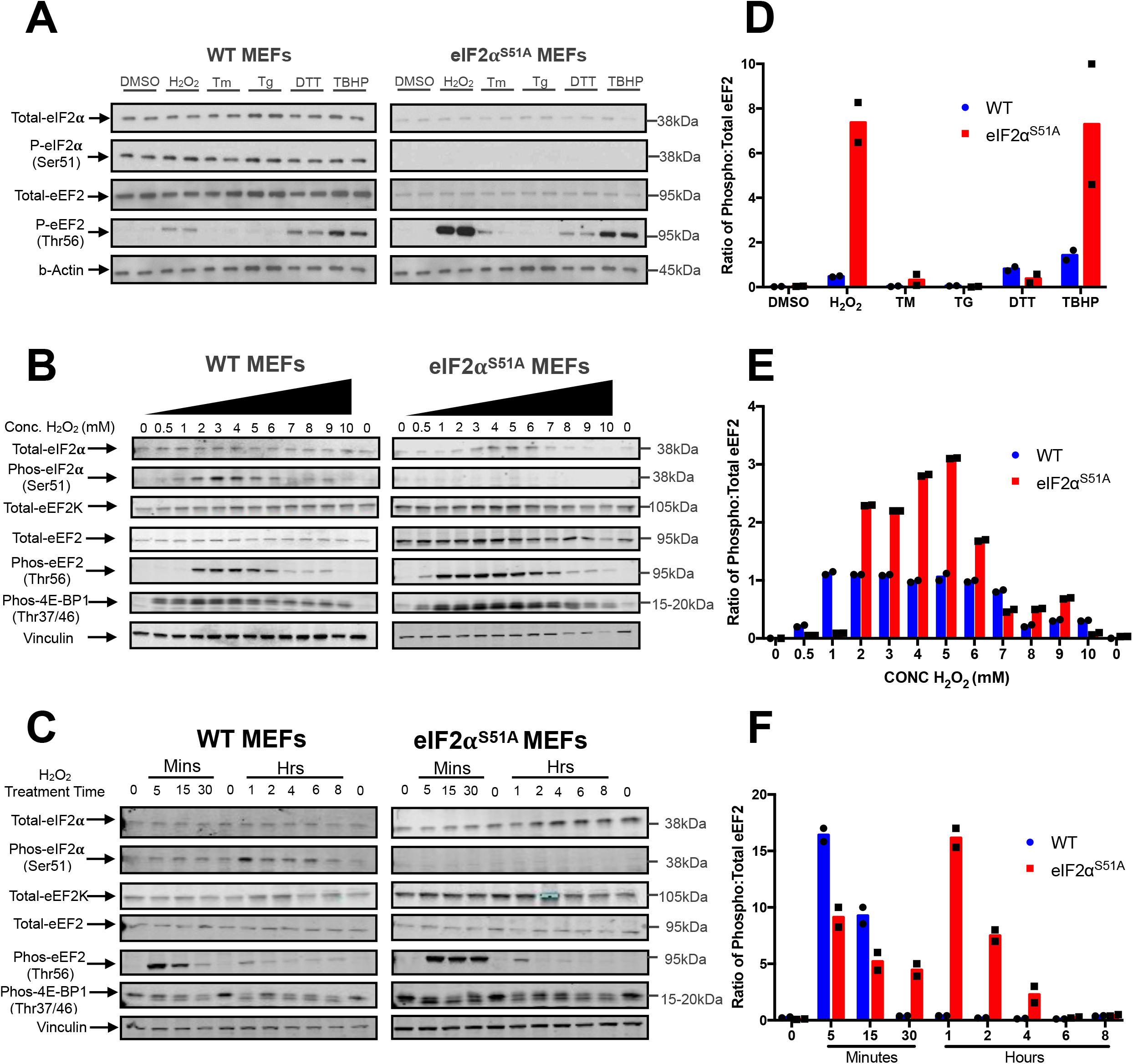
eEF2 phosphorylation under oxidative stress. (**A**) Western blot analysis was performed with lysates from wildtype and eIF2α^S51A^ mutant MEFs. Cells were treated with the same agents as indicated in Figure 2C. Cells were treated for 1 hour except in the case of tunicamycin (Tm), which was added for 4 hours. (**B**) Western blot analysis was performed with lysates from wildtype and eIF2α^S51A^ mutant MEFs treated with increasing doses of H_2_O_2_ (0.5-10 mM) for 1 hour. (**C**) Western blot analysis was performed with lysates from wildtype and eIF2α^S51A^ mutant MEFs treated with 500 μM H_2_O_2_ for increasing periods (5 min-8 hr). (**D**) Quantification of eEF2 phosphorylation. The graph represents the ratio of phosphorylated to total eEF2 quantified from the Western blots in Figure 5A. The intensities were quantified using Licor Image Studio software. The individual symbols represent individual data points. The individual data points represent duplicated individual repeat experiments. (**E**) Quantification of eEF2 phosphorylation. The graph represents the ratio of phosphorylated to total eEF2 quantified from the Western blots in Figure 5B. The intensities were quantified using Licor Image Studio software. The individual symbols represent individual data points. The individual data points represent duplicated individual repeat experiments. (**F**) Quantification of eEF2 phosphorylation. The graph represents the ratio of phosphorylated to total eEF2 quantified from the Western blots in Figure 5C. The intensities were quantified using Licor Image Studio software. The individual symbols represent individual data points. The individual data points represent duplicated individual repeat experiments.

Dose-response studies revealed marked differences in the sensitivity of eEF2 phosphorylation to H_2_O_2_. Whereas eEF2 phosphorylation in wildtype MEFs peaked between 2 – 4 mM H_2_O_2_ and coincided with eIF2α phosphorylation, eEF2 phosphorylation was observed with as little as 0.5 mM H_2_O_2_ in eIF2α^S51A^ MEFs (Figure 5B). In both MEF lines, eEF2 phosphorylation subsided at high doses of H_2_O_2_, likely due to cytotoxicity (Figure 6B,C). No major cell line difference was observed in the sensitivity to H_2_O_2_-mediated changes in the phosphorylation of 4E-binding protein 1 (4E-BP1), which increased at low doses and subsided at high doses (Figure 5B). The finding that 4E-BP1 phosphorylation increased rather than decreased in response to H_2_O_2_ suggested that inhibition of TORC1 signaling is unlikely to be responsible for inhibiting translation under oxidative stress. The role of the paradoxical increase in 4E-BP1 phosphorylation in response to H_2_O_2_, which would be expected to promote eIF4E-eIF4G interaction and translation (Chu et al., 2016) remains presently unclear.

**Figure 6:**
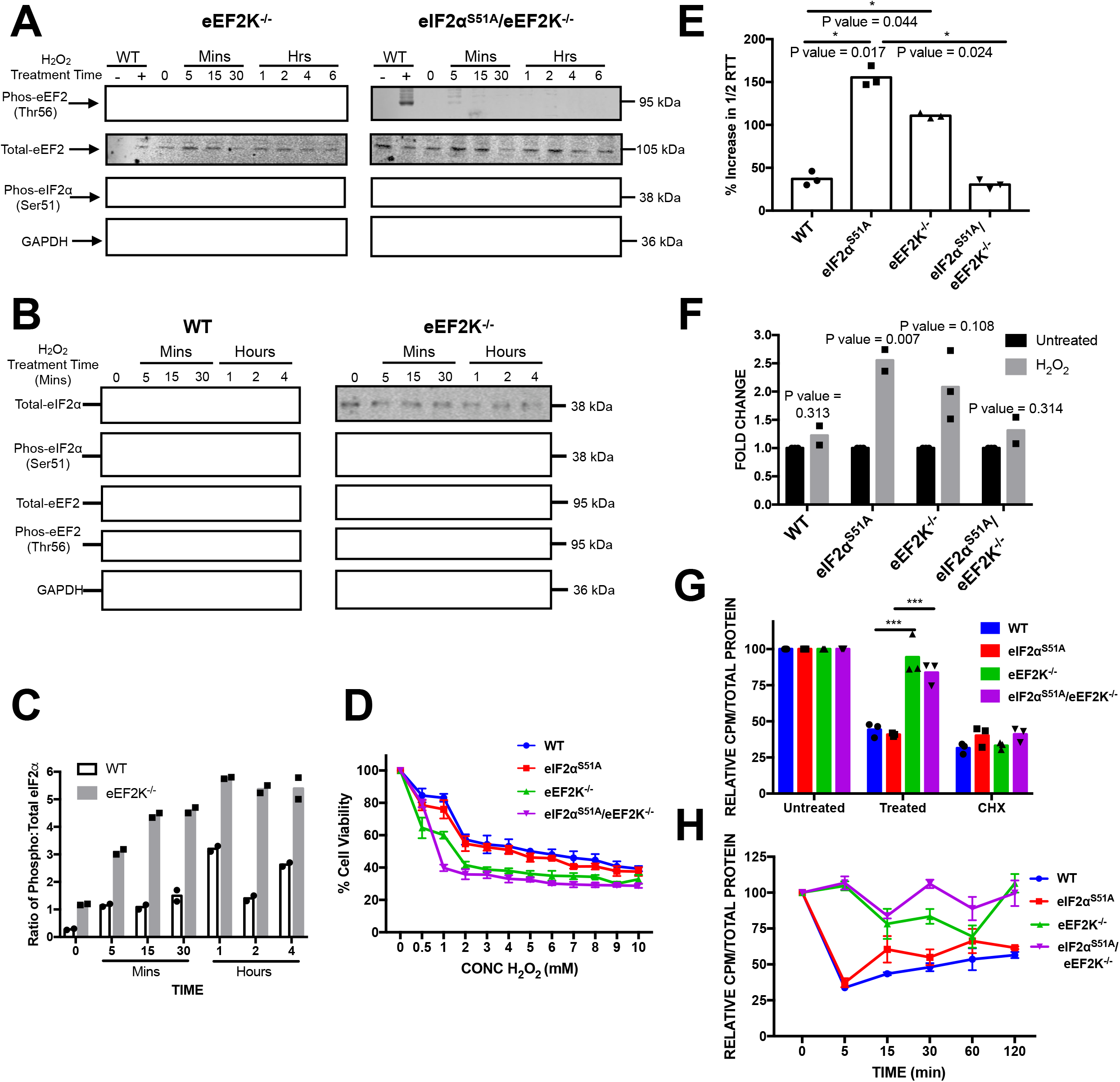
Role of eEF2K in eIF2α-independent translational inhibition. (**A**) Western blot analysis was performed with lysate prepared from eEF2K^−/−^ and eIF2α^S51A^/eEF2K^−/−^ MEFs treated with 500 μM H_2_O_2_ for increasing periods (5 min - 6 hr). Untreated WT MEFs were used as a negative control and WT MEFs treated with 500 μM H_2_O_2_ for 5 minutes were used as a positive control. (**B**) Wildtype and eEF2K^−/−^ MEFs were exposed to 500 μM H_2_O_2_ for increasing periods (5 min - 4 hr), and eIF2α phosphorylation was assessed by immunoblotting. (**C**) Quantification of eIF2α phosphorylation. The graph represents the ratio of phosphorylated to total eIF2α quantified from the Western blots in Figure 6B. The intensities were quantified using Licor Image Studio software. The individual symbols represent individual data points. The individual data points represent duplicated individual repeat experiments. (**D**) Cells were treated with the indicated doses (0.5 - 10 mM) of H_2_O_2_ for 1 hour. A MTT assay was performed to assess cell viability. The graph represents the means ± standard deviations of 8 replicates. (**E**) Ribosome transit times were analyzed in wildtype, eIF2α^S51A^, eEF2K^−/−^ and eIF2α^S51A^/eEF2K^−/−^ MEFs treated with 500 μM H_2_O_2_. The graph represents the mean percent increases in 1/2 transit times ± standard deviations of three independent experiments each done in technical triplicates. The individual symbols represent summaries of the individual experiments. *P value between WT and eIF2α^S51A^ = 0.017, P value between WT and eEF2K^−/−^ = 0.044, P value between eIF2α^S51A^ and eEF2K^−/−^ = 0.024. (**F**) The graph represents the normalized mean fold change of 1/2 transit times ± standard deviations of the same experiments as shown in Figure 6D. The individual symbols represent summaries of the individual experiments. *P value WT = 0.313, P value eIF2α^S51A^ = 0.007, P value eEF2K^−/−^ = 0.108, P value eIF2α^S51A^/eEF2K^−/−^ = 0.314. (**G**) Cells were either treated with 1 mM H_2_O_2_, 50 μg/mL CHX or untreated. After 15 minutes, [^35^S]-methionine was added for an additional 45 minutes. Protein synthesis levels were then quantified and adjusted according to the total amount of protein. The graph represents normalized means ± standard deviations of at two independent experiments each done in technical triplicates. The individual symbols represent summaries of the individual experiments. *P value eEF2K^−/−^ = 0.004, P value eIF2α^S51A^/eEF2K^−/−^ = 0.0007. (**H**) Cells were treated with 1 mM H_2_O_2_ for increasing times indicated and labeled with [^35^S]-methionine for 5 minutes. Protein synthesis levels were then quantified and adjusted according to the total amount of protein. The graph represents the normalized means ± standard deviations of at least two independent experiments each done in technical triplicates.

Time course experiments also revealed a marked difference in the kinetics of eEF2 phosphorylation. Whereas phosphorylation was induced within 5 minutes of H_2_O_2_ exposure in both MEF lines, the response was more sustained in eIF2α^S51A^ MEFs (Figure 5C). H_2_O_2_-induced eIF2α phosphorylation in wildtype MEFs, which we determined to be mediated, at least to a large extent, by PERK (Figure S4), did not peak until 1 hour after stress application, indicating that it may have a function in maintaining the translation bock for increased periods. In summary, this data shows that eEF2 phosphorylation occurs in an acute yet transient manner under oxidative stress. Conversely, eIF2α phosphorylation is delayed but more sustained.

### eEF2K knockout mouse embryonic fibroblasts are deficient in attenuating translation under oxidative stress

To establish a possible functional role of eEF2K in eIF2α-independent translation regulation under oxidative stress, eEF2K was knocked out using the CRISPR-Cas9 system in wildtype and eIF2α^S51A^ MEFs. When challenged with 500 μM H_2_O_2_ for 5 minutes to 6 hours, neither cell line exhibited eEF2 phosphorylation (Figure 6A). Comparing wildtype and eEF2K^−/−^ MEFs, we found that basal and H_2_O_2_-induced eIF2α phosphorylation is considerable augmented in cells lacking eEF2K (Figure 6B,C). Treatment of eEF2K^+/+^ and eEF2K^−/−^ MEFs with increasing concentrations of H_2_O_2_ (0.5 - 10 mM for 1 hour) revealed increased sensitivity of both eEF2K knockout lines to H_2_O_2_-induced cytotoxicity relative to their parental lines with the eIF2α^S51A^ eEF2K^−/−^ MEFs being the most sensitive (Figure 6D). The ~2.5-fold increase in ribosome transit time observed in eIF2α^S51A^ MEFs treated with H_2_O_2_, was reduced to ~2-fold in eEF2K^−/−^ and less than 1.5-fold in eEF2K^−/−^/eIF2α^S51A^ MEFs (Figure 6E, F). Continuous metabolic labeling during a 1 hour challenge with H_2_O_2_, revealed that eEF2K^−/−^ MEFs are unable to attenuate protein synthesis to the same extent as the parental wildtype and eIF2α^S51A^ MEFs (Figure 6G). The same result was obtained in 5-minute pulse labeling experiments across a 2 hour time course of H_2_O_2_ challenge (Figure 6H). Taken together, these data show that eEF2K is required for initiating translational inhibition under oxidative stress.

### eEF2K is necessary for inhibiting translation under oxidative stress in *S. pombe*

To investigate whether eEF2K-mediated mechanism of translation inhibition is evolutionarily conserved, the role of eEF2K in protein synthesis under stress was studied in *S. pombe.* The *cmk2* gene, encoding the putative S. pombe orthologue of eEF2K, was deleted in wildtype and eIF2α^S52A^ mutant strains. By quantitative phosphoproteomics, we identified a 2.12 (+/-0.3)-fold increase in the phosphorylation of a putative MAPK site, serine 436, of Cmk2p after 15 minutes of exposure to H_2_O_2_, indicating that Cmk2p phosphorylation is H_2_O_2_-inducible (Figure S6E). In addition, loss of *cmk2* confers sensitivity to H_2_O_2_ (Sánchez-Piris et al., 2002).

Acute exposure to H_2_O_2_ revealed hypersensitivity of *Δcmk2* and *eIF2α^S52A^ Δcmk2* strains (Figure 7A). Both the *WT* and *eIF2α^S52A^* strains showed decreased levels of protein synthesis comparable to the cycloheximide control (Figure 7B). The *Δcmk2* and *eIF2α^S52A^ Δcmk2* strains, however, exhibited significantly higher levels of translation compared to the wildtype and eIF2α^S52A^ strains (Figure 7B). Protein synthesis was also measured by short pulse labeling with [^35^S]-methionine at various times after H_2_O_2_ (5 mins – 2 hrs). After 5 minutes, *WT* and *eIF2α^S52A^* strains showed a >50% reduction in protein synthesis, whereas *Δcmk2* and *eIF2α^S52A^ Δcmk2* strains showed only a ~25% reduction (Figure 7C). The same differential effect was seen at the 2-hour time point when *Δcmk2* and *eIF2α^S52A^ Δcmk2* cells had partially recovered protein synthesis. Unlike MEFs deficient in eEF2K, *S. pombe* cells lacking *cmk2* still downregulated protein synthesis between 15 and 60 minutes after H_2_O_2_ challenge (Figure 7C). It is thus possible that a partially redundant eEF2 kinase exists in fission yeast, perhaps the calmodulin-dependent kinase, Cmk1p. Regardless, in both S. pombe and mammalian cells, eEF2K facilitates acute inhibition of translation in response to oxidative stress.

**Figure 7:**
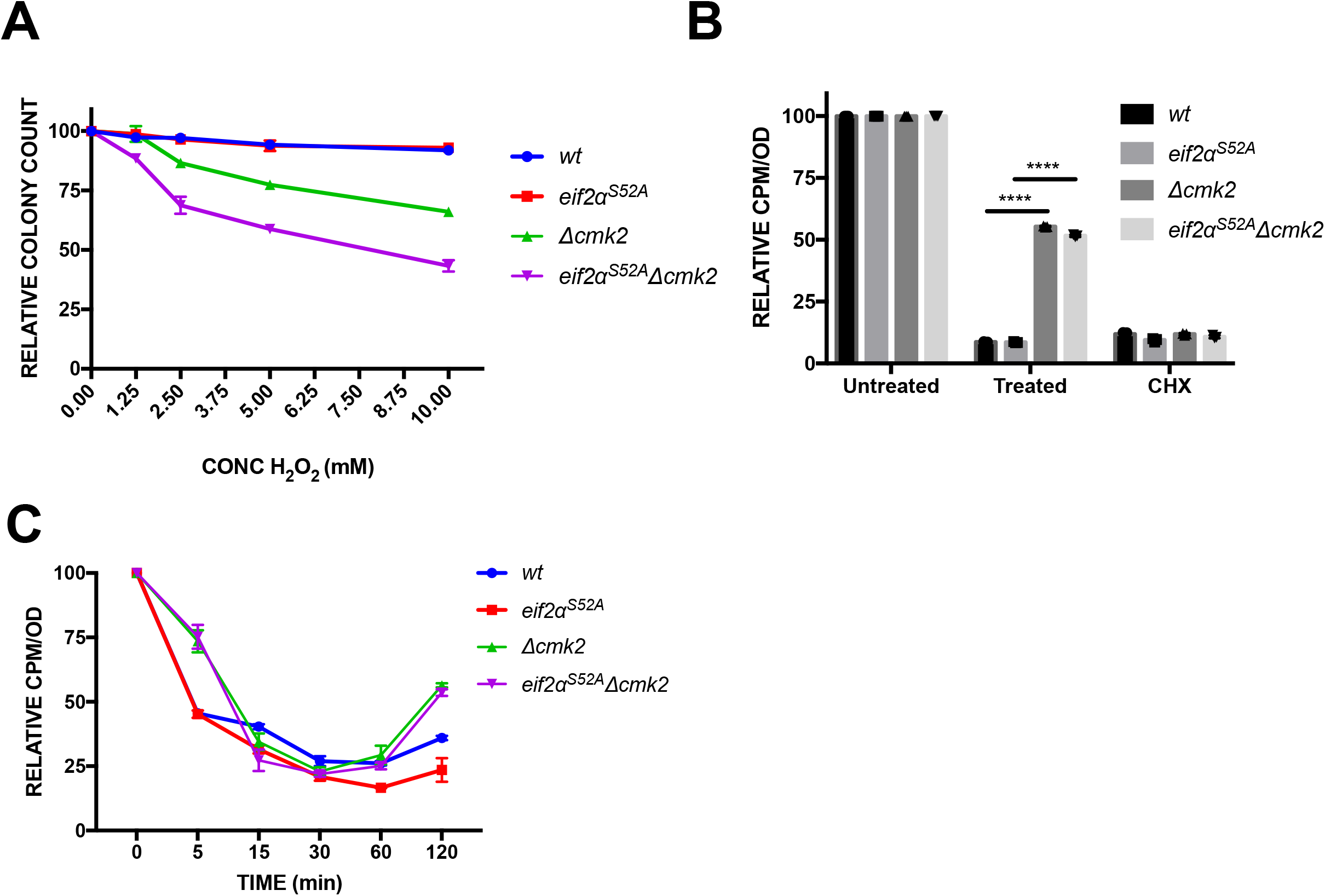
Effect of hydrogen peroxide on viability and protein synthesis in *S. pombe cmk2* mutants. (**A**) Cells were treated with increasing concentrations (0 – 1 mM) of H_2_O_2_ for 1 hour. An equal number of cells were plated, grown for 3 days, and colonies were counted. The graph represents the mean ± standard deviations of at least two independent experiments each done in triplicates. (**B**) Cells were either treated with 1 mM H_2_O_2_, 50 μg/mL CHX or untreated. After 15 minutes, [^35^S]-methionine was added for an additional 45 minutes. Protein synthesis levels were quantified and adjusted according to the total amount of protein. The graph represents the normalized mean ± standard deviations of two independent experiments each done in triplicates. The individual symbols represent summaries of the individual experiments. *P value eEF2K^−/−^ = 1.2 x 10^−9^, P value eIF2α^S51A^/eEF2K^−/−^ = 9.4 x 10^−9^. (**C**) Cells were treated with 1 mM H_2_O_2_ for the periods indicated and labeled with [^35^S]-methionine for 5 minutes. Protein synthesis was quantified and normalized to the total amount of protein. The graph represents the normalized mean ± standard deviations of at least two independent experiments each done in triplicates.

## DISCUSSION

### Significance of blocking translation elongation in response to oxidative stress

The main observation of the present study is that, unlike with ER stress, acute translational shutdown in response to oxidative stress occurs independently of eIF2α phosphorylation despite robust eIF2α kinase pathway activation. Rather, we demonstrate in MEFs that oxidative stress triggers rapid activation of eEF2K and inhibitory phosphorylation of eEF2 thus effecting a block in elongation. While most known mechanisms of translational control are exerted at the level of initiation, more recently, control at the elongation step has come into focus (Richter and Coller, 2015). For example, genome-wide ribosome profiling in budding yeast has revealed that oxidative stress blocks elongation, especially in the first 50 codons (Gerashchenko et al., 2012; Wu et al., 2019). A slowdown of elongation was also described in E. coli under hyperosmotic stress (Dai et al., 2018). Together with our observations in MEFs, these findings suggest that a block in translation elongation is an evolutionarily conserved primary response to oxidative stress that may also be invoked under other stress conditions.

### Temporally distinct roles of eIF2α and eEF2 phosphorylation in MEFs

If eIF2α is dispensable for translational shutdown, does it have any role in the response to oxidative stress? Our genetic data indicate that yeast and mammalian cells deficient in both, eIF2α and eEF2 phosphorylation, are supersensitive to killing by H_2_O_2_ (Figure 6, 7). It thus appears that both pathways cooperate in the defense against oxidative stress. Our kinetic data in MEFs show that eEF2 phosphorylation is a rapid and transient event (~ 5 – 15 minutes), whereas eIF2α phosphorylation peaks with a delay but is sustained (~ 1 – 6h). This suggests a temporal ordering of both events with eEF2 triggering the initial block in protein synthesis and eIF2α phosphorylation maintaining it (Figure 8A). This conclusion is supported by our demonstration that cells deficient in eEF2 phosphorylation fail to initiate inhibition of protein synthesis, whereas MEFs deficient in eIF2α phosphorylation recover from the arrest precociously.

**Figure 8:**
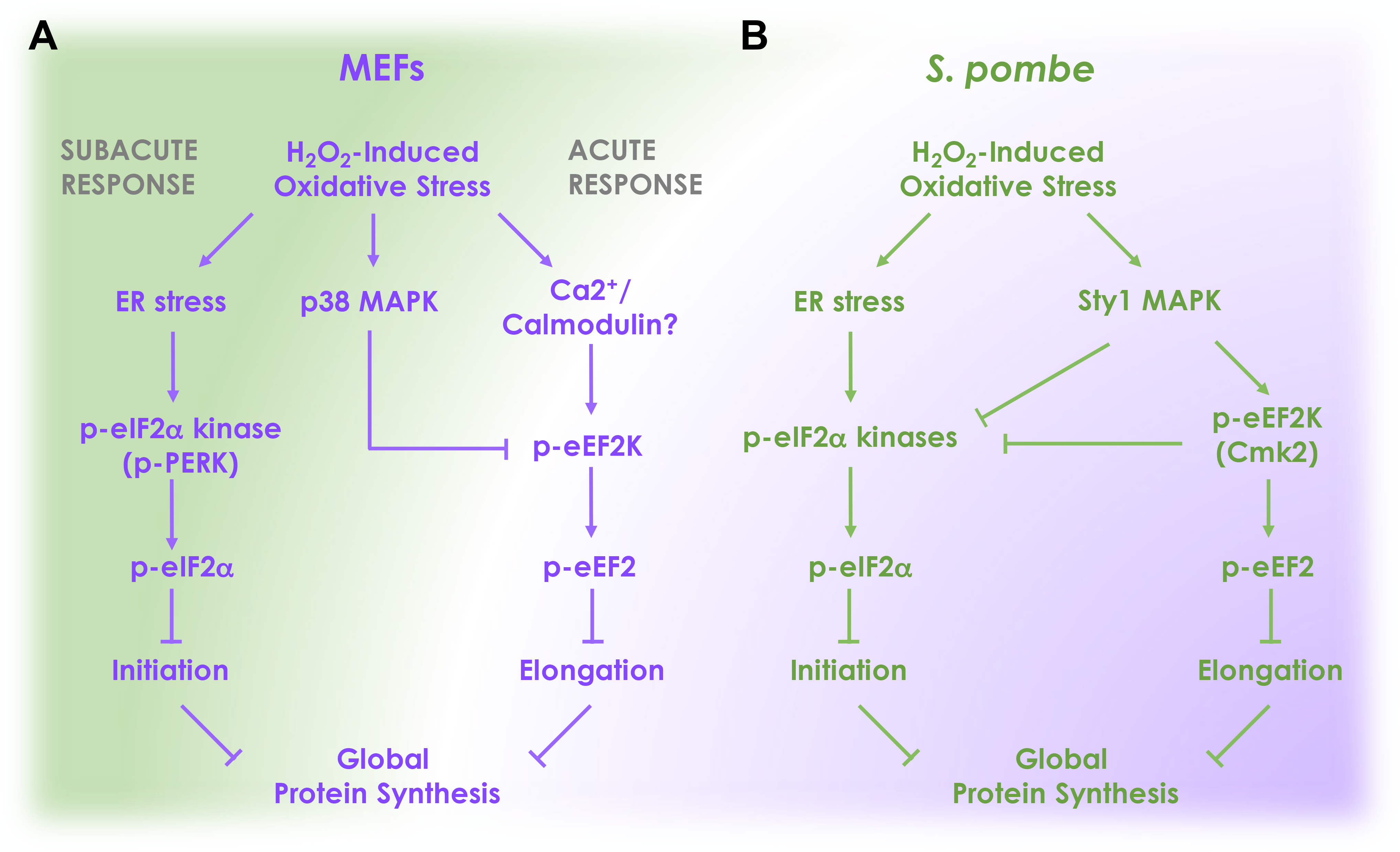
Model of the conserved and divergent aspects of stress-induced translation inhibition pathways in MEFs and S. pombe. (**A**) Model for MEFs. Oxidative stress triggers two parallel pathways. Rapid activation of eEF2K leads to acute arrest of translation elongation. The accumulation of misfolded proteins in the endoplasmic reticulum caused by oxidative stress triggers PERK-dependent eIF2α phosphorylation and subacute block in translation elongation. Signaling through p38 MAPK creates a negative feedback loop to terminate eEF2K signaling. (**B**) Model for S. pombe. Oxidative stress rapidly induces both eIF2α phosphorylation and eEF2K activation. Sty1p MAPK activation triggers eEF2K signaling and creates a negative feedback loop to terminate eIF2α phosphorylation. There is also evidence that Cmk2p suppresses eIF2α phosphorylation.

The reliance on the instant arrest in protein synthesis afforded by blocking elongation as compared to the delayed response achieved by a block in initiation suggests that the initial translational stress response is primarily geared toward limiting the accumulation of damaged nascent proteins, whereas the delayed response at the level of initiation may serve resource conservation purposes. Arresting ribosomes may also enable expedient resumption of elongation upon passing a putative quality control step thus being economically advantageous.

The timing difference in the eIF2α and eEF2 phosphorylation responses may be explained by distinct upstream triggers, a rapid kinase cascade in the case of eEF2K/eEF2, and the delayed accumulation of unfolded proteins in the case of PERK/eIF2α. The exact nature of the kinase cascade culminating in eEF2K activation remains unknown and may differ between stress conditions and organisms.

### Conserved versus organism-specific response patterns

Whereas eIF2α phosphorylation is dispensable for H_2_O_2_-induced translational shutdown in both MEFs and S. pombe due to a block in translation elongation, the responses also show marked organism-specific differences rendering the model in Figure 8A inapplicable to S. pombe. First, the kinetics of eIF2α phosphorylation are much faster in S. pombe than in MEFs with strong induction seen as early as 5 minutes after exposure to H_2_O_2_ (Berlanga et al., 2010; Dunand-Sauthier et al., 2005). Thus, phosphorylation of eIF2α and eEF2 appear to be initiated in parallel rather than in succession. Since H_2_O_2_-induced eIF2α phosphorylation lasts for at least 120 minutes (Krohn et al., 2008), it could theoretically play a role in maintaining the translation block. However, our data suggest that, in S. pombe, Cmk2p rather than eIF2α phosphorylation is required for maintenance of translational inhibition two hours after H_2_O_2_ treatment (Figure 7C). This is consistent with data showing that Cmk2p attenuates eIF2α phosphorylation (Sanchez-Marinas et al., 2018).

A second marked difference between MEFs and S. pombe is in upstream signaling. The MAPK Sty1p is known to directly phosphorylate Cmk2p in response to H_2_O_2_ in S. pombe (Sánchez-Piris et al., 2002). At the same time, Sty1p activation limits eIF2α phosphorylation thus creating a negative feedback loop to terminate the translation arrest (Berlanga et al., 2010; Dunand-Sauthier et al., 2005) (Figure 8B). In contrast, while mammalian p38 MAPK kinase activity is also increased by H_2_O_2_ (Figure S5A), this kinase is known to inhibit rather than activate eEF2K (Knebel et al., 2002). Confirming this, we have found that MAPK inhibitors potently prolong H_2_O_2_-induced eEF2 phosphorylation in MEFs (Figure S5B). It thus appears that, unlike in S. pombe, oxidative stress activates eEF2K in a MAPK-independent fashion in mammalian cells. Since H_2_O_2_ increases intracellular calcium within seconds (Hu et al., 1998; Jornot et al., 1999), eEF2K may be directly activated by calmodulin in response to oxidative stress. Activation of p38 may subsequently act to terminate eEF2K activity for translation elongation to resume (Figure 8A).

### Crosstalk between eIF2α and eEF2 phosphorylation in MEFs

The pattern of eEF2 phosphorylation in MEFs is markedly disturbed by the absence of eIF2α phosphorylation, indicating crosstalk between the two translation regulatory pathways. Not only is eEF2 phosphorylation triggered at lower doses of H_2_O_2_ (0.5 vs. 2 mM), the response is also substantially prolonged (2 hr versus 30 min). Thus, the pathway blocking elongation is hypersensitized and hyperactivated in the absence of the pathway inhibiting initiation. The hyperactivation of eEF2 phosphorylation suggests an eIF2α phosphorylation-dependent sensing mechanism that activates a negative feedback loop to inhibit eEF2, perhaps mediated by p38 kinase (Figure 8A). The hypersensitization of eEF2 phosphorylation in cells deficient in eIF2α phosphorylation might be explained by an increased load of reactive oxygen species or reduced antioxidant capacity in eIF2α^S51A^ mutant MEFs as indicated by the stress profile apparent in the RNAseq data at baseline.

The pattern of eIF2α phosphorylation was also disturbed in cells lacking eEF2K, showing hyperactivation by H_2_O_2_ (Figure 6B,C). Likewise, in S. pombe, eIF2α phosphorylation in response to oxidative stress is prolonged in *cmk2* deleted cells (Sanchez-Marinas et al., 2018), suggesting that eIF2α phosphorylation is intensified. This could simply be due to increased accumulation of misfolded proteins in cells deficient in acute elongation arrest of protein synthesis, although more intricate crosstalk scenarios are also conceivable.

### Implications for cancer

Within the tumor microenvironment, cancer cells are frequently exposed to oxidative stress which arises from oncogenic stimulation, metabolic defects, hypoxia and nutrient depletion (Gorrini et al., 2013). The ability to respond to oxidative stress is thus central to the survival of cancer cells. The pro-survival function attributed to the frequent overexpression of eEF2K in cancer has been proposed to be due to improved stress adaptation (Fu et al., 2014). Nutrient depletion, for example, leads to AMPK-dependent activation of eEF2K and a block in elongation, whereas loss of eEF2K hampers the growth of tumors in mice under caloric restriction (Leprivier et al., 2013). High eEF2K expression correlates with decreased overall survival in medulloblastoma (Leprivier et al., 2013) and, as our own data mining shows, in renal cancer (Figure S7). The data presented here demonstrate that eEF2K activation is also a rapid consequence of exposure to oxidative stress. Considering that many clinically used cancer therapeutics act by exerting oxidative stress, the status of the eEF2K pathway as well as its crosstalk with the eIF2α phosphorylation pathway may set a threshold that determines therapeutic response. Both pathways, possibly in combination, therefore appear attractive cancer drug targets, although effective eEF2K inhibitors remain to be developed (Liu and Proud, 2016).

## LIMITATIONS OF THE STUDY

One limitation of the present study is the use of WT and eIF2α^S51A^ mutant MEFs after long-term adaptation to tissue culture. While these two cell lines are supposed to vary in only one amino acid residue in eIF2α, our mRNA expression profiles show many differences that point to more substantial divergence of the two MEF lines. Acute, inducible knockout of the eIF2α^S51A^ phosphorylation site would thus be preferable but no such system is currently available. We have mitigated this deficiency by performing additional experiments in S. pombe cells, which support our main conclusions that eIF2α phosphorylation is dispensable for translational arrest in response to H_2_O_2_, a response that is rather mediated by a block in translation elongation through eEF2K activation. A second limitation is that we observed higher than desirable variability in the response of eIF2α phosphorylation to H_2_O_2_. Whereas peak induction was consistently observed after ~1 hour of H_2_O_2_ treatment, early time points showed greater variability. This might be due to slight but difficult to control variations in the stress status of individual MEF cultures as we sometimes observed considerable baseline activation of eIF2α phosphorylation in the absence of H_2_O_2_. While this led to inconsistent quantifications at early time points, this limitation does not impact the main conclusions of our study. A final limitation is that we present tantalizing evidence of cross-talk between eIF2α phosphorylation and eEF2 phosphorylation but the molecular mechanism mediating this cross-talk remains to be determined in future studies.

## Supporting information

Downregulated Mutant 120 Metascape Files

Upregulated Mutant 120 Metascape Files

Downregulated Mutant 15 Metascape Files

Upregulated Mutant 15 Metascape Files

Downregulated WT 120 Metascape Files

Upregulated WT 120 Metascape Files

Downregulated WT 15 Metascape Files

Upregulated WT 15 Metascape Files

Upregulated Time 0 Metascape Files

Downregulated Time 0 Metascape Files

## AUTHOR CONTRIBUTIONS

Conceptualization, M.S. and D.A.W.; Methodology, M.S., A.R.C., C.-C.Y., P.A.-B., and P.M.; Formal Analysis, M.S. and A.R.C.; Investigation, M.S., Y.-Y. L., C.-C. Y., A.R.C., P.M., and D.A.W.; Writing – Original Draft, M.S. and D.A.W.; Writing – Review & Editing, M.S., Y.-Y. L., C.-C. Y., A.R.C., P.M., and D.A.W.; Visualization, M.S.; Supervision, D.A.W. and P.A.-B.; Funding Acquisition, M.S. and D.A.W.

## ACKNOWLEDGEMENTS

This research was funded by grants R21 CA190588, R01 GM105802, and R01 GM121834 (D.A.W.) and an F31 fellowship CA210616-01 (M.S.) from the U.S. National Institutes of Health (NIH). Part of this work was funded by NIH P30 grants CA030199 and GM085764. D.A.W. is a scholar of the 1000 Talent Program funded by the Government of the People’s Republic of China. We thank Beata Grallert and Erik Boye for providing the *eIF2α^S52A^* S. pombe strain, César de Haro for the *Δhri1 Δhri2 Δgcn2* strain, and Randal Kaufman for the eIF2α^S51A^ MEFs, syngeneic wildtype MEFs, and discussion.

