## Supplemental Information for "Crosstalk between eIF2α and eEF2 phosphorylation pathways optimizes translational arrest in response to oxidative stress"

### Transparent Methods

#### Reagents

Reagents are listed in Table S2.

#### Cell Lines and Tissue Culture

*Wildtype and eIF2α$^{S51A}$ Mutant Mouse Embryonic Fibroblasts (MEFs)*

Both wildtype and eIF2α$^{S51A}$ mutant mouse embryonic fibroblast cell lines were generously provided by Randal J. Kaufman. MEF cell lines were cultured in DMEM containing 4.5 mg/mL glucose (Invitrogen), 2.0 mM glutamine, 10% fetal bovine serum and 1% penicillin-streptomycin. Cells were incubated in 5% $CO_2$ at 37°C.

*Generation of eEF2K Knockout and eEF2K$^{-/-}$/eIF2α$^{S51A}$ MEFs*

Wildtype and eIF2α$^{S51A}$ mutant MEFs were transduced with a CRISPR/Cas9 lentiviral system containing pools of sgRNA libraries targeting the mouse eEF2K gene. Cells were transduced for 48 hours. Single cells were then sorted to generate monoclonal populations. Monoclonal cell lines were tested for functional eEF2K by Western blotting. To confirm eEF2K knockout, cellular DNA was isolated and genetic deletion was confirmed by PCR. Cell lines were also sequenced to further confirm eEF2K knockout in both wildtype and eIF2α$^{S51A}$ mutant MEFs.

#### *S. pombe* Strains

*Wildtype and eIF2α$^{S52A}$ Mutant Strains*

Strains were maintained in rich yeast extract medium with supplements (YES) or Edinburgh minimal media (EMM) at 30°C. Growth was measured using optical density at 596 nm. The genotypes of the strains used in this study are outlined in Table S1.

*Generation of Δcmk2 and eIF2α$^{S52A}$ Δcmk2 Strains*

The deletion of *cmk2* was generated in wildtype and *eIF2α$^{S52A}$* mutant strains by PCR amplification of the plasmid pfa6a-NAT-HA. Cells were then transformed, selected on nourseothricin (NAT) plates, and *cmk2* knockout was confirmed by PCR.

#### MTT Cell Viability Assay

Cells were treated with indicated treatments or DMSO alone (0.1% final concentration) for indicated times. Viability was assessed by MTT assay (ATCC®) using the manufacturer's protocol. Briefly, 10 µl of MTT reagent was added to cells for 2 hrs. 100 µl of detergent reagent was then added to cells and the absorbance was read at 570 nm. Results are averages of 8 replicates ± standard deviations.

#### Protein Extraction and Western Blot Analysis

*Mouse Embryonic Fibroblast Sample Preparation*

For protein extraction, mammalian cells were washed once with 1x phosphate-buffered saline (PBS), scraped off and lysed in RIPA lysis buffer (50 mM Tris-HCl pH 8.0, 150 mM NaCl, 0.1% NP-40, 0.1% SDS and 12 mM sodium deoxycholate) supplemented with Halt™ Protease Inhibitor Cocktail (Thermo Scientific) and incubated on ice for 15 mins followed by sonication. After a 10 min centrifugation (13,000 rpm at 4°C), supernatants were taken for protein quantification following Pierce™ BCA Protein Assay Kit (Thermo Scientific). Samples were boiled with SDS sample buffer. Equal amounts of protein (20-40 µg) were resolved on 4-20% tris-glycine SDS–PAGE gels and transferred to nitrocellulose membranes, which were blocked in 5% bovine serum albumin (BSA) in 1x tris-buffered saline (TBS) (10 mM Tris-HCl pH 7.5, 150 mM NaCl) for 1 hr and then incubated with 5% BSA in TBST (10 mM Tris-HCl pH 7.5, 150 mM NaCl and 0.1% Tween 20) containing primary antibody overnight at 4°C. Membranes were washed (TBST 3×) and incubated in TBST containing LICOR fluorescent secondary antibody. After washing (TBST 3×), immunoreactive bands were detected using the LICOR system.

*S. pombe Sample Preparation*
Yeast cells were cultured and collected in exponential growth phase (OD 0.5 – 1.0). Cells were centrifuged and resuspended in cold RIPA lysis buffer supplemented with Halt™ Protease Inhibitor Cocktail (Thermo Scientific) and added to 600 µl of zirconia/silica beads (BioSpec). Cells were ruptured in a FastPrep instrument for 40 sec at level 6. When necessary, supernatants were taken for protein quantification following Pierce™ BCA Protein Assay Kit (Thermo Scientific). Lysates were diluted with 2x SDS sample buffer and denatured at 95°C for 10 mins before loading. Western blot analysis was performed.

**Protein Synthesis Fluorescence Assay**
Protein synthesis levels were assessed in a non-radioactive manner by fluorescent protein synthesis assay (Protein Synthesis Assay Kit, Cayman Chemical) using the manufacturer's protocol. Briefly, corresponding treatments were added to cells for indicated times in a 96 well plate. Cycloheximide (50 µg/mL) was added for 1 hr as a positive control for protein synthesis inhibition. Cells were then incubated with O-propargyl-puromcyin (2.5 µg/mL) followed by fixation, washing (3x), and staining with 5 FAM-azide. Cells were then washed (3x) and fluorescence was detected with a fluorescent plate reader (excitation/emission = 485/535).

**[$^{35}$S]-Methionine Labeling**
*Mouse Embryonic Fibroblast Sample Preparation*
Cells were plated in 10 cm dishes at 3 × $10^6$ cells/dish in supplemented DMEM, incubated overnight and treated with 500 µM $H_2O_2$ for indicated times. Cycloheximide (50 µg/mL) was added for 1 hr as a control for protein synthesis inhibition. Cells were then labeled with 10 µCi/mL [$^{35}$S]-methionine (EasyTag™ EXPRESS$^{35}$S Protein Labeling Mix, Perkin Elmer) for 5 mins, rinsed once with cold 1x PBS and harvested immediately by scraping. Cells were pelleted and resuspended in 0.5 mL lysis buffer (25 mM Tris-HCl pH 8.0, 150 mM NaCl, 0.1% NP-40, 0.1% SDS and 12 mM sodium deoxycholate) supplemented with Halt™ Protease Inhibitor Cocktail (Thermo Scientific). Lysates were boiled at 95°C for 10 mins, briefly centrifuged and blotted on Whatman filter paper for total protein precipitation. Filters were precipitated with ice-cold 20% TCA for 10 mins followed by 10% TCA for 5 mins. Filters were rinsed twice with 100% ethanol for 5 mins and air-dried. Radioactivity levels were determined by liquid scintillation counting. Total cellular protein was quantified with Pierce™ BCA Protein Assay Kit (Thermo Scientific) following the manufacturer's instruction. Incorporation of [$^{35}$S] methionine into total cellular protein was calculated and plotted.

*S. pombe Sample Preparation*
Yeast samples were centrifuged, resuspended in RIPA lysis buffer and added to 600 µl of zirconia/silica beads (BioSpec). Cells were ruptured in a FastPrep instrument for 40 sec at level 6. Lysates were diluted with 2x SDS sample buffer and denatured at 95°C for 10 mins before blotting.

**Polysome Profile Analysis**
*Mouse Embryonic Fibroblast Sample Preparation*
Cells (4 × $10^6$) were seeded in 15 cm culture dishes and grown to ~70% confluence. Following treatment with 500 µM $H_2O_2$ at indicated time points, cycloheximide (CHX) (100 µg/mL) was added to cells for 5 min at 37°C. Cells were washed twice with cold 1x PBS containing CHX (100 µg/mL), scraped gently in 5 mL of ice cold 1x PBS containing CHX (100 µg/mL). Cells were centrifuged at 200 × g for 5 min at 4°C. The cell pellets were suspended in 0.45 mL of polysome lysis buffer (5 mM Tris-HCl at pH 7.5, 2.5 mM $MgCl_2$, 1.5 mM KCl). Lysis buffer was supplemented with 5 µl of 10 mg/mL CHX, 1 µl of 1 mM DTT, 100 units RNase inhibitor (RNaseOUT, Invitrogen), 1x Pierce™ protease inhibitor and 1x Pierce™ phosphatase inhibitor. Cells were vortexed for 5 sec followed by the addition of 25 µl 10% sodium deoxycholate and 25 µl Triton-X 100. Lysates were then vortexed again for 5 sec and centrifuged at 16,000 × g for 7 min at 4°C. Supernatants (cytosolic cell extracts) were collected and measured in absorbance of 260 nm. Approximately 10-15 ODs of lysates were layered over 5%–50% cold sucrose gradients in buffer (200 mM HEPES-KOH at pH 7.4, 50 mM MgCl2, 1 mM KCl, 100 µg/mL CHX and 1x

Pierce™ protease inhibitor). Gradients were centrifuged at 39,000 rpm in a Beckman SW28 rotor for 2 hr at 4°C. After centrifugation, 14 equal-sized fractions (0.75 mL/fraction) were collected and analyzed through UV detection.

*S. pombe Sample Preparation*
Cells in exponential growth phase (OD 0.5 – 1.0) were treated with CHX (100 µg/mL) and centrifuged for 5 min at 2500 × g at 4°C. Cells were lysed in polysome lysis buffer (20 mM Tris-HCl [pH 7.5], 50 mM KCl, 10 mM $MgCl_2$, and 1 mM DTT) supplemented with 100 µg/mL CHX and 1x Pierce™ protease inhibitor. Lysates were added to 600 µl of zirconia/silica beads (BioSpec) and ruptured in a FastPrep instrument for 20 sec at level 6. Supernatants were extracted through centrifugation and polysome profiles were obtained.

**Ribosome Half-Transit Time Measurement**
Ribosome transit time refers to the length of time needed for a ribosome, after attaching to a mRNA, to complete translation and release a finished polypeptide. This is measured by analyzing the kinetic flow of radioactivity from polysome-bound (nascent) polypeptides to completed (released) polypeptides. Nascent and released polypeptides are separated by differential centrifugation. Since at any one time there is, on the average, one-half of a completed polypeptide per ribosome on a mRNA molecule, determination of the kinetics of flow of radioactivity as described will yield one half-transit time values. Measurements of half-transit times are completely independent of rates of attachment of ribosomes to mRNA and of the number of polyribosomes (Fan and Penman, 1970; Gehrke et al., 1981).

Cells ($3 \times 10^6$) were plated in 10 cm dishes in supplemented DMEM and incubated overnight. Cells were simultaneously labeled with 10 µCi/mL [$^{35}$S]-methionine (EasyTag™ EXPRESS$^{35}$S Protein Labeling Mix, Perkin Elmer) and treated with 500 µM $H_2O_2$. At the times indicated, cells were washed with cold 1x PBS containing 100 µg/mL CHX and harvested immediately. Cells were pelleted and resuspended in 0.45 mL of lysis buffer (10 mM Tris-HCl at pH 7.5, 15 mM $MgCl_2$, 10 mM NaCl, 100 µg/mL CHX) supplemented with Halt™ Protease Inhibitor Cocktail (Thermo Scientific). Cells were lysed by adding 25 µl of 10% Triton X-100 and 25 µl 10% sodium deoxycholate, followed by vortexing for 5 seconds. Nuclei and mitochondria were pelleted by centrifugation for 15 min at maximum speed in a microfuge at 4°C. 200 µl of the post-mitochondrial supernatant (PMS) was saved to measure [$^{35}$S]-methionine incorporation into total protein (nascent and completed proteins). Ribosomes were pelleted by centrifugation of the remaining 200 µl of the PMS at 90,000 × g for 1 hr at 4°C in a Beckman TLA120 rotor. 200 µl of the post-ribosomal supernatant (PRS) were removed to measure the incorporation of [$^{35}$S]-methionine into completed protein. 50 µl of PMS and PRS samples from indicated time points were precipitated with TCA after spotting on Whatman filter paper. Filters were washed with ice-cold 20% TCA for 10 mins followed by 10% TCA for 5 mins. Filters were then rinsed twice with 100% ethanol for 5 mins and air-dried before being subjected to liquid scintillation counting.

**RNA Sequencing**
Total RNA from cells untreated or treated with 500 µM $H_2O_2$ at times 0, 15 minutes and 120 minutes was obtained using the RNeasy Mini Kit (Qiagen) following the manufacturer's instructions. RNA quality was assessed using a Bioanalyzer (Bio-Rad Experion), and RNA sequencing was performed at the Genomics Facility (Sanford Burnham Prebys Medical Discovery Institute) on the Illumina platform. The averaged expression data for ~10,000 mRNAs was narrowed down to ~1,600 mRNAs with significant differences in summed FPKM values of transcripts, p value ≤ 0.05. For each condition, these mRNAs were hierarchically clustered using Spearman Rank Correlation and Complete Linkage Clustering. A list of 613 mRNAs specific to WT MEFs that were significantly regulated (≥ 3-fold, p ≤ 0.05) by $H_2O_2$ was imported into Metascape (www.metascape.org) and canonical pathways enriched in the datasets were identified for all conditions in both cell lines (Figure 1B). A list of 982 mRNAs specific to eIF2α$^{S51A}$ MEFs that were significantly regulated (≥ 3-fold, p value ≤ 0.05) by $H_2O_2$ was imported into Metascape and canonical pathways enriched in the dataset were identified for all conditions in both cell lines (Figure 1C). The complete dataset is provided in Table S3 and available in the Gene Expression Omnibus (accession number PRJNA517725).

**Statistical Analysis**
Statistical analyses of replicate datasets were performed with Graphpad Prism. Typically, data were averaged, standard deviations calculated, and statistical significance was assessed using the T Test assuming two-tailed distribution and unequal variance.

**Poly-A Fragment Sequencing**
RNA sequencing was performed by the Genomics Core Facility at Sanford Burnham Prebys Medical Discovery Institute under the direction of Brian James. PolyA RNA was isolated using the NEBNext® Poly(A) mRNA Magnetic Isolation Module and barcoded libraries were made using the NEBNext® Ultra II™ Directional RNA Library Prep Kit for Illumina®(NEB, Ipswich MA). Libraries were pooled and single end sequenced (1X75) on the Illumina NextSeq 500 using the High output V2 kit (Illumina Inc., San Diego CA). Read data was processed in BaseSpace (basespace.illumina.com). Reads were aligned to to Mus musculus genome (mm10) using STAR aligner (https://code.google.com/p/rna-star/) with default settings. Differential transcript expression was determined using the Cufflinks Cuffdiff package (https://github.com/cole-trapnell-lab/cufflinks).

**Metascape Pathway Enrichment Analyses (metascape.org)**
User-provided gene identifiers are first converted into their corresponding M. musculus Entrez gene IDs using the latest version of the database (last updated on 2017-03-16). If multiple identifiers correspond to the same Entrez gene ID, they will be considered as a single Entrez gene ID in downstream analyses. For each given gene list, pathway and process enrichment analysis has been carried out with the following ontology sources: GO Biological Processes, KEGG Pathway, Reactome Gene Sets and CORUM. All genes in the genome have been used as the enrichment background. Terms with a p-value < 0.01, a minimum count of 3, and an enrichment factor > 1.5 (the enrichment factor is the ratio between the observed counts and the counts expected by chance) are collected and grouped into clusters based on their membership similarities. More specifically, p-values are calculated based on the accumulative hypergeometric distribution, and q-values are calculated using the Benjamini-Hochberg procedure to account for multiple hypotheses. Kappa scores are used as the similarity metric when performing hierachical clustering on the enriched terms, and sub-trees with a similarity of > 0.3 are considered a cluster. The most statistically significant term within a cluster is chosen to represent the cluster.

**Western Blot Data Quantification**
Licor imaging software was used to quantify Western blot band intensities. To quantify a given band, the total amount of signal detected in the area that contains the band is expressed as the sum of the intensities measured in all of these pixels. The signal value represents an unbiased estimate of specific signal intensity and is not affected by adjustments to the image display. A portion of this signal, however, corresponds to background, due to dark current, membrane reflection, autofluorescence, non-specific antibody binding, etc. Therefore, to estimate the amount of signal due only to the specific binding of the antibodies, a background subtraction is performed. To perform background subtraction, a background area surrounding the band is first defined. Within this area the average intensity of each pixel is calculated as an estimation of non-specific signal in the vicinity of the band of interest. This value is subtracted from the intensity of each pixel within the band area. The reported Signal value corresponds to the sum of the residual intensity values (after background subtraction) of all of the pixels in the band.

**FACS Analysis**
MEFs were either untreated or treated with 500 μM $H_2O_2$ for 2 hours. Cells were trypsinized and resuspended in media without FBS. Cells were treated with 20 μM 2',7'–dichlorofluorescin diacetate (DCFDA) for 30 minutes. Cells were then resuspended in cold 1 x PBS. FACS analysis was performed by the Flow Cytometry Core at Sanford Burnham Prebys Medical Discovery Institute using the LSRFortessa.

**Determination of doubling time of *S. pombe* cultures**

*S. pombe* strains were inoculated into liquid YES media at an OD260 between 0.015 and 0.03 and grown for 24 hours. OD was measured at time 0, 5, 10, 21 and 23 hours and an exponential growth curve was fitted. The time to doubling the OD from 0.5 to 1 was calculated by solving the exponential equations for y -0.5 and y = 1.

**Cell Counting Assay**
Cells were treated with the indicated treatments for 1 hour, trypsinized and resuspended in media. Cells were diluted 1:2 in trypan blue and the number and concentration of viable cells was calculated using the Nexcelcom Bioscience Cellometer Auto T4 Bright Field Cell Counter and corresponding software.

**Polysome Run-off by Glucose Withdrawal**
In yeast, the action of glucose as a signaling molecule affects a diverse number of biochemical pathways. It has been shown that glucose depletion by means of glucose withdrawal from the growth medium led to a rapid, almost complete inhibition of protein synthesis. Re-addition of glucose causes a rapid reversal of this inhibition. The inhibition does not come about via a gross decay of mRNA, and neither the inhibition nor its reversal by re-addition of glucose requires transcription of new mRNAs (Ashe et al., 2000). The “runoff” of polysomes observed after glucose removal requires that translational elongation continues while initiation is inhibited (Mathews et al., 1996). Translational elongation of a polypeptide requires at least two GTP molecules per amino acid added, whereas initiation requires only one or two GTP molecules per polypeptide chain (Merrick and Hershey, 1996). It was also concluded glucose deprivation an effective way to block translation initiation independently of eIF2α phosphorylation (Ashe et al., 2000).

Cells in exponential growth phase (OD 0.5 – 1.0) were pelleted and resuspended in yeast extract medium without glucose and incubated at 30°C for 5 min. Cells were then with treated with CHX (100 µg/mL) and centrifuged for 5 min at 2500 × g at 4°C. Cells were lysed in polysome lysis buffer (20 mM Tris-HCl [pH 7.5], 50 mM KCl, 10 mM $MgCl_2$, and 1 mM DTT) supplemented with 100 µg/mL CHX and 1x Pierce™ protease inhibitor. Lysates were added to 600 µl of zirconia/silica beads (BioSpec) and ruptured in a FastPrep instrument for 20 sec at level 6. Supernatants were extracted through centrifugation and polysome profiles were obtained.

**Supplemental Data Items**
**Table S1 related to Transparent Methods: *S. pombe* Strains**

| Name | Genotype | Used in Fig. |
|---|---|---|
| WT | h+ ade6-M210 ura4-d18 | 3A, 3B, 3C, 3D, 4D, 6A, 7B, 7C, S3 |
| eIF2α$^{S52A}$ | h+ ade6-M210 eIF2alpha-S52A::ura4-d18 | 3A, 3B, 3C, 3D. 4D, 7A, 7B, 7C, |
| Δcmk2 | h+ ade6-M210 ura4-d18 cmk2::NAT | 7A, 7B, 7C, |
| eIF2α$^{S52A}$ Δcmk2 | h+ ade6-M210 eIF2alpha-S52A::ura4-D18 cmk2::NAT | 7A, 7B, 7C, |
| Δhri1 Δhri2 Δgcn2 | h- ade6- 216 leu1-32 ura4-d18 his7-366 hri1::ura4 hri2::leu1 gcn2::ura4 | 3A, 3B, 3C, 3D |
| Δsty1 | h- leu1-32 ura4-d18 sty1::ura4 | 3C, 3D |

**Table S2 related to Transparent Methods: Reagents**

| Reagent | Company | Cat. No. | Storage | Dilution/Conc |
|---|---|---|---|---|
| Hydrogen Peroxide | Sigma Aldrich | H1009-100mL | -20°C | 0.1 – 1 mM |
| Tert-butyl hydroperoxide | Acros Organics | 75-91-2 | -20°C | 200 µM |
| Cycloheximide | Acros Organics | 66-81-9 | -20°C | 50 µg/mL |
| Tunicamycin | TOCRIS | 351610 | -20°C | 10 µg/mL |
| Thapsigargin | TOCRIS | 11381 | -20°C | 1 – 10 µM |

| | | | | |
|---|---|---|---|---|
| DMSO | TOCRIS | 3176 | RT | |
| Dithiothreitol | Sigma Aldrich | 3483-12-3 | -20°C | 10 mM |
| Anti-Total eIF2α | Cell Signaling | 9722S | -20°C | 1:1000 |
| Anti- Phospho eIF2α (Ser51) | Cell Signaling | 9721S | -20°C | 1:500 |
| Anti- Total eEF2 | Cell Signaling | 2332S | -20°C | 1:1000 |
| Anti- Phospho eEF2 (Thr56) | Cell Signaling | 2331S | -20°C | 1:500 |
| Anti- GAPDH | GeneTex | GTX100118 | -20°C | 1:1000 |
| Anti- b-Actin | GeneTex | GTX109639 | -20°C | 1:1000 |
| Anti- Total eEF2K | Novus | NBP1-51329 | -20°C | 1:500 |
| Anti- Phospho 4E-BP1 (Thr37/46) | Cell Signaling | 2855P | -20°C | 1:500 |
| Anti- Vinculin | Sigma Aldrich | V4505-100UL | -20°C | 1:1000 |
| Anti- Total p38 | Cell Signaling | 9212 | -20°C | 1:1000 |
| Anti- Phospho p38 (Thr180/Tyr182) | Cell Signaling | 4631S | -20°C | 1:500 |
| Bovine Serum Albumin | Alpha Diagnostic Inc. | 80400-100 | -20°C | 5% |
| Donkey anti Rabbit 680CW Secondary | LICOR | 925-68073 | -20°C, 4°C | 1:10,000 |
| Goat anti Mouse 680CW Secondary | LICOR | 925-68070 | -20°C, 4°C | 1:10,000 |
| Goat anti Rabbit 800CW Secondary | LICOR | 925-32211 | -20°C, 4°C | 1:10,000 |
| Goat anti Mouse 800CW Secondary | LICOR | 925-32210 | -20°C, 4°C | 1:10,000 |

**Table S3 related to Figure 1: RNA Sequencing Gene Lists**
**Table S4 related to Figure 1: Metascape Gene Lists**

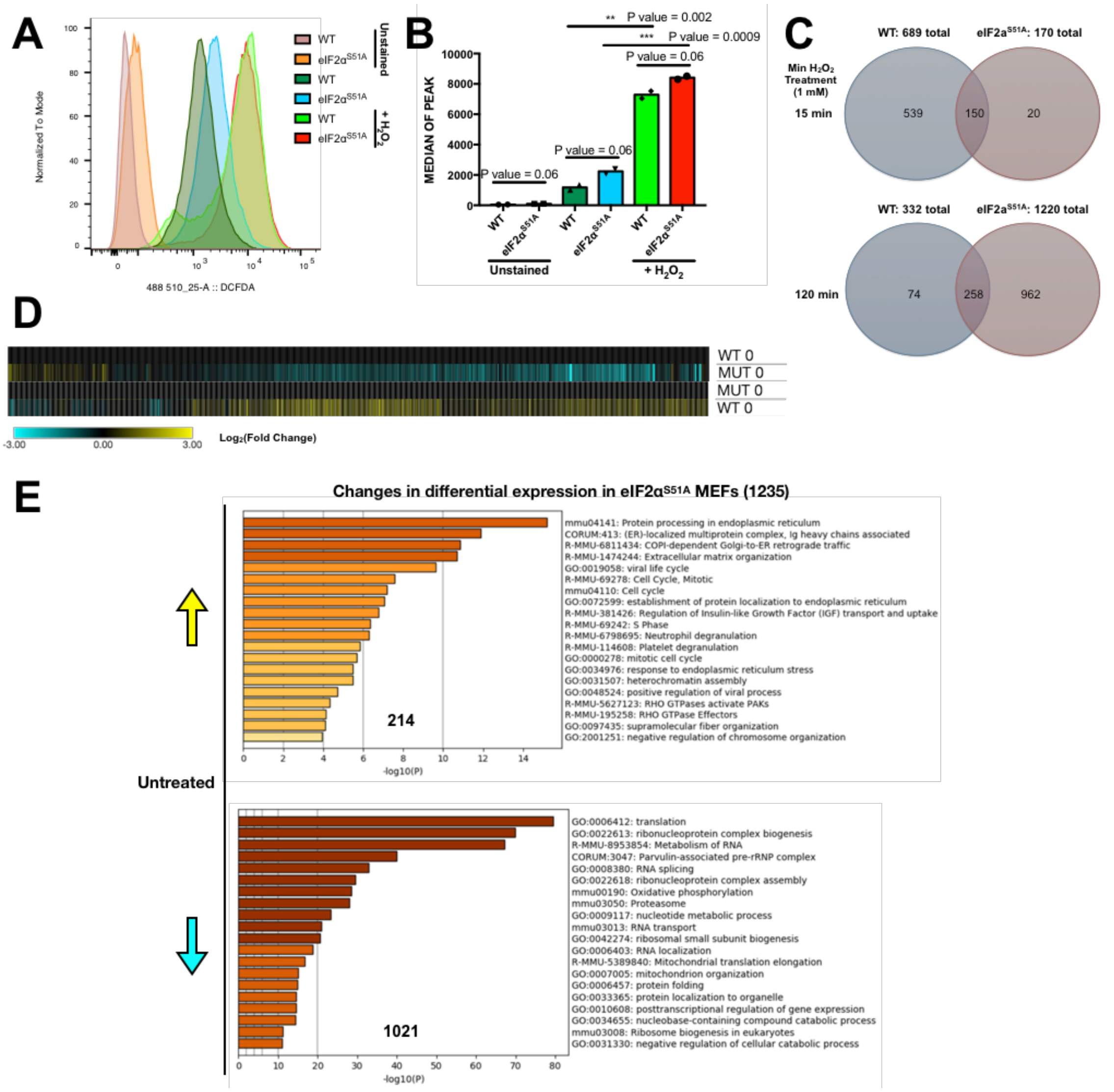


**Figure S1. $H_2O_2$-induced changes in ROS and mRNA levels, Related to Figure 1.**
(**A**) The effects of $H_2O_2$ on ROS levels. Cells were either untreated or treated with 500 μM $H_2O_2$ for 2 hrs. Cells were collected and stained with DCFDA for 30 minutes and FACS analysis was performed.
(**B**) The graph represents a duplicates, each individual symbol represents an individual data point. There is a fold change of 1.2 between the medians of WT and eIF2α$^{S51A}$ MEFs treated with $H_2O_2$ and a fold change of 1.8 between the medians of untreated WT and eIF2α$^{S51A}$ MEFs.
(**C**) Significant $H_2O_2$-induced changes in mRNA levels ($p \leq 0.05$) at both 15 and 120 minutes of $H_2O_2$ treatment (1 mM) were identified. There were 613 total changes occurring only in wildtype MEFs between both time points and 982 occurring only in eIF2α$^{S51A}$ mutant MEFs at both time points. The total number of changes at each time point for each cell line include the overlap.
(**D**) Total RNA was extracted from WT and and analyzed by RNAseq. The data represents a compilation of significant changes in differential expression ($p \leq 0.05$) found in untreated WT and at baseline. Results are normalized to either untreated WT MEFs or untreated eIF2α$^{S51A}$ MEFs at time 0 as indicated and displayed as a clustered heat map. Upregulated transcripts are shown in yellow and downregulated transcripts in blue.
(**E**) Individual lists of induced and repressed mRNAs (±3-fold change, $p \leq 0.05$) in eIF2α$^{S51A}$ MEFs compared to WT MEFs were loaded into Metascape and the pathways enriched are indicated at time 0. Colored boxes represent either upregulated (yellow) or downregulated (blue) pathways.

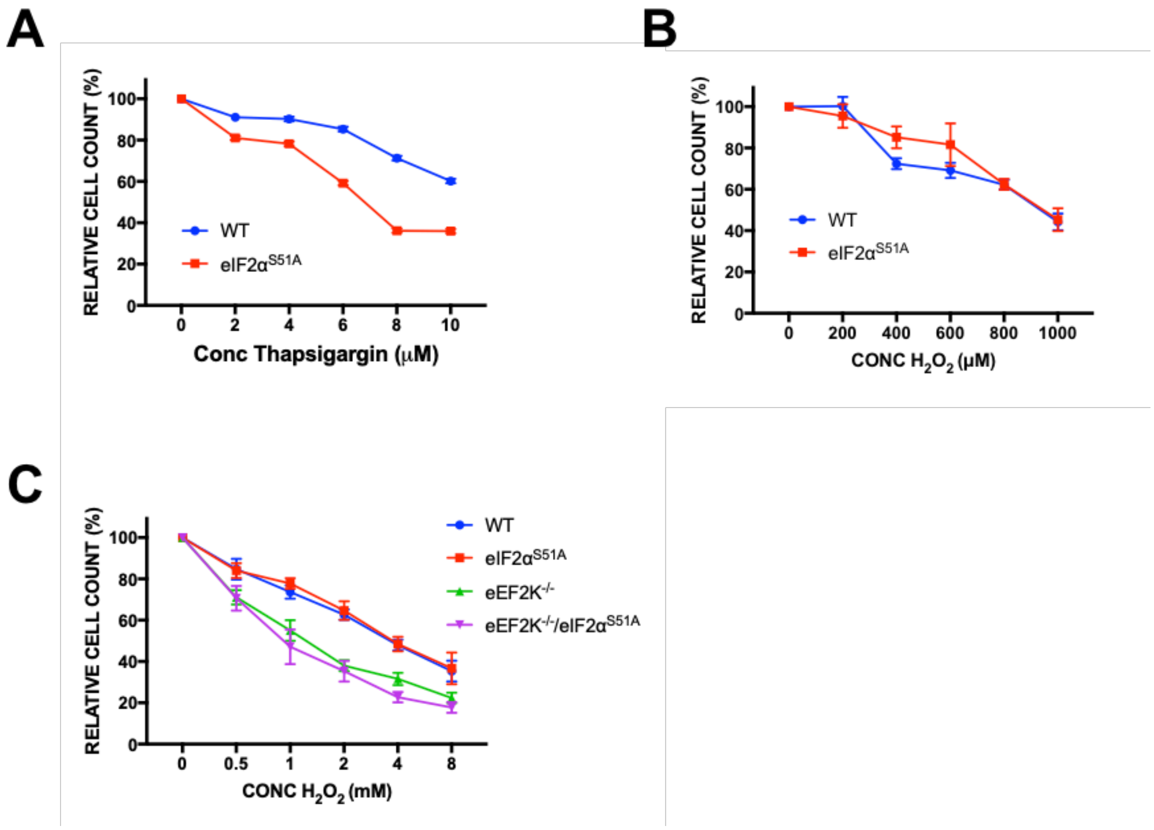


**Figure S2. The effects of ER and oxidative stress on cell viability, Related to Figure 2 and Figure 6.**

(**A**) MTT assay results in Figure 2A were validated by cell counting assay. The graph represents a summary of 3 individual experiments each performed in duplicates.

(**B**) MTT assay results in Figure 2B were validated by cell counting assay. The graph represents a summary of 3 individual experiments each performed in duplicates.

(**C**) MTT assay results in Figure 6B were validated by cell counting assay. The graph represents a summary of 3 individual experiments each performed in duplicates.

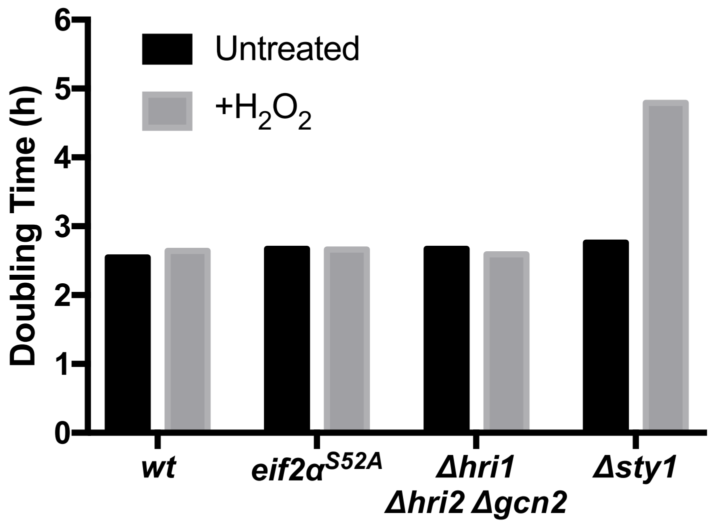


**Figure S3. The effects of $H_2O_2$ on doubling time in *S. pombe*, Related to Figure 3.**
S. pombe strains were treated with 1 mM $H_2O_2$ and grown for 24 hours. Doubling times were calculated based on OD measured at various time points.

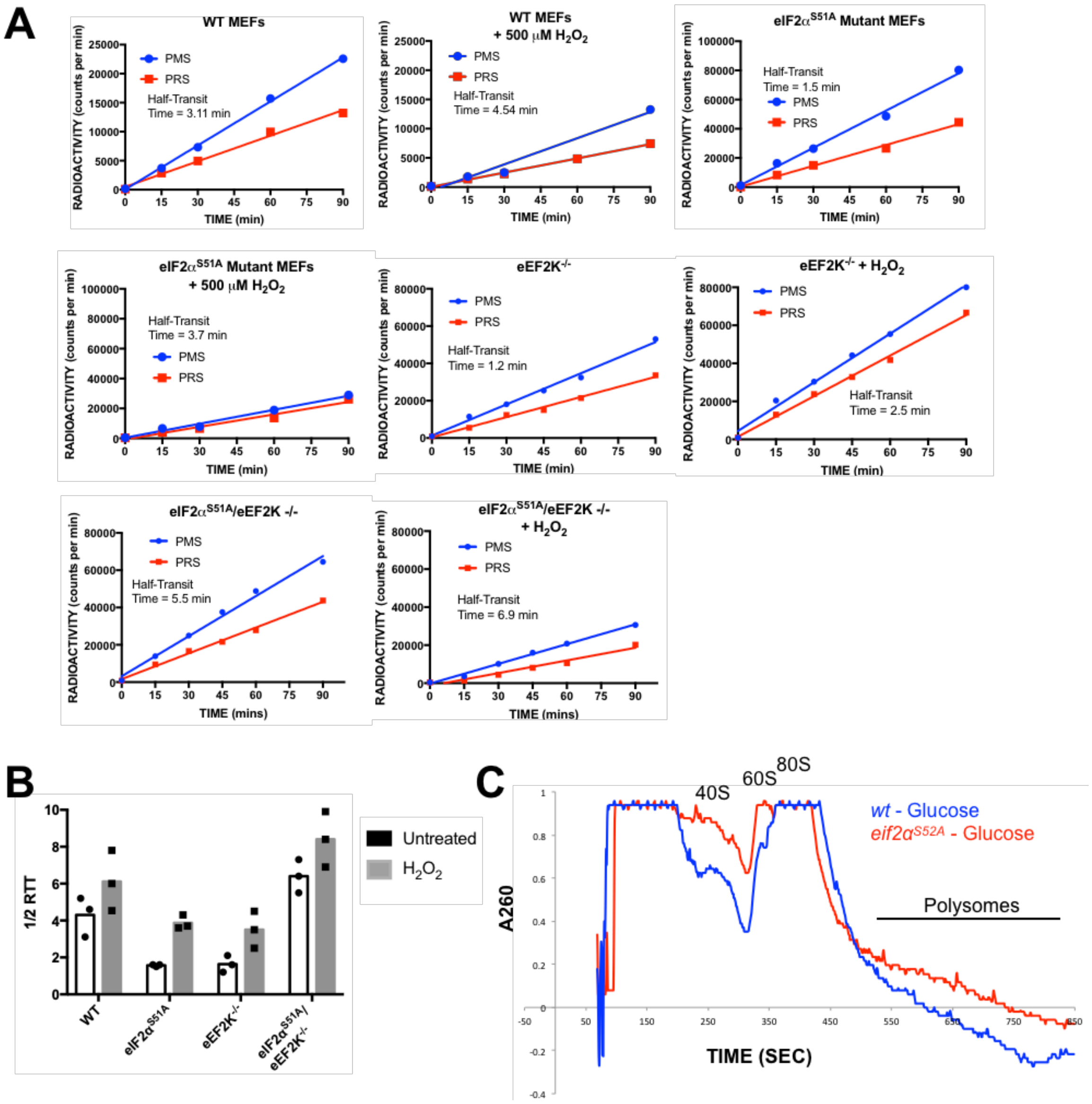

**Figure S4. Additional elongation and initiation analysis, Related to Figure 4.**
(**A**) Ribosome transit time experiments. The ribosome half-transit times of WT, eIF2α$^{S51A}$, eEF2K$^{-/-}$ and eIF2α$^{S51A}$/eEF2K$^{-/-}$ MEFs treated with and without 500 μM $H_2O_2$ were determined as described in Materials and Methods. Incorporation rates of [$^{35}$S]-methionine into total protein within the PMS and PRS was obtained by linear regression analysis, which was used to calculate half-transit times. Each experiment was performed three separate times.
(**B**) The graph represents a summary of each individual replicate experiment that was performed and the half-transit times (min) to obtain the results and statistical analysis represented in Figure 4B, 4C, 6C and 6D.
(**C**) Polysome profiles of *S. pombe* strains under glucose deprivation. *Wt* and *eIF2α$^{S52A}$* strains were deprived of glucose for 5 minutes and polysome profiles were acquired.

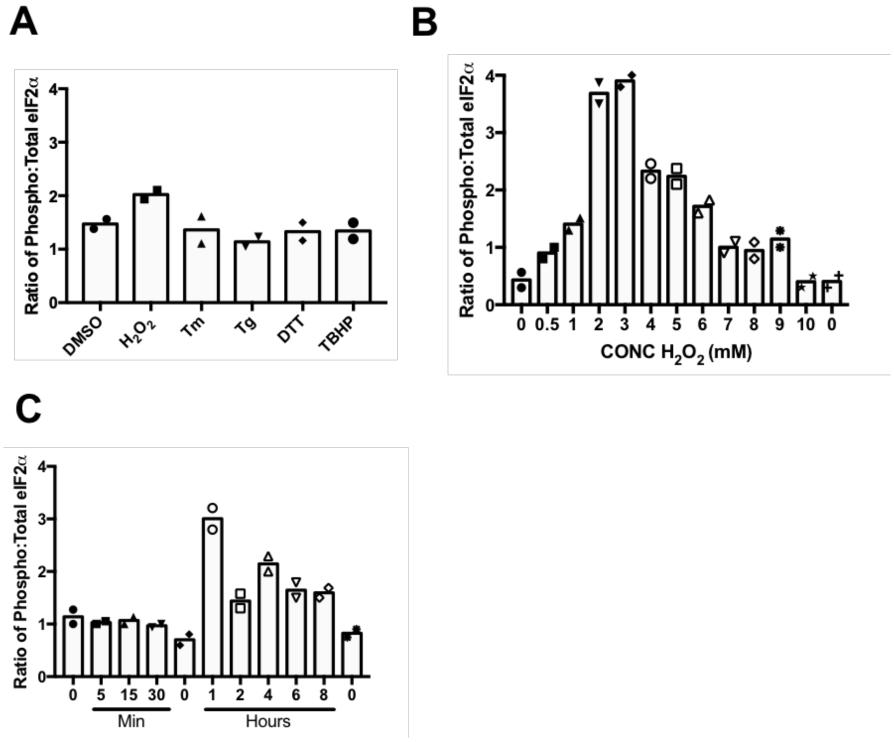


**Figure S5. Quantification of eIF2α phosphorylation, Related to Figure 5.**
(**A**) Quantification of eIF2α phosphorylation. The graph represents the ratio of phosphorylated to total eIF2α quantified from the Western blots representing WT MEFS in Figure 5A. The intensities were quantified using Licor Image Studio software. The individual symbols represent individual data points. The individual data points represent duplicated individual repeat experiments.
(**B**) Quantification of eIF2α phosphorylation. The graph represents the ratio of phosphorylated to total eIF2α quantified from the Western blots representing WT MEFS in Figure 5B. The intensities were quantified using Licor Image Studio software. The individual symbols represent individual data points. The individual data points represent duplicated individual repeat experiments.
(**C**) Quantification of eIF2α phosphorylation. The graph represents the ratio of phosphorylated to total eIF2α quantified from the Western blots representing WT MEFS in Figure 5C. The intensities were quantified using Licor Image Studio software. The individual symbols represent individual data points. The individual data points represent duplicated individual repeat experiments.

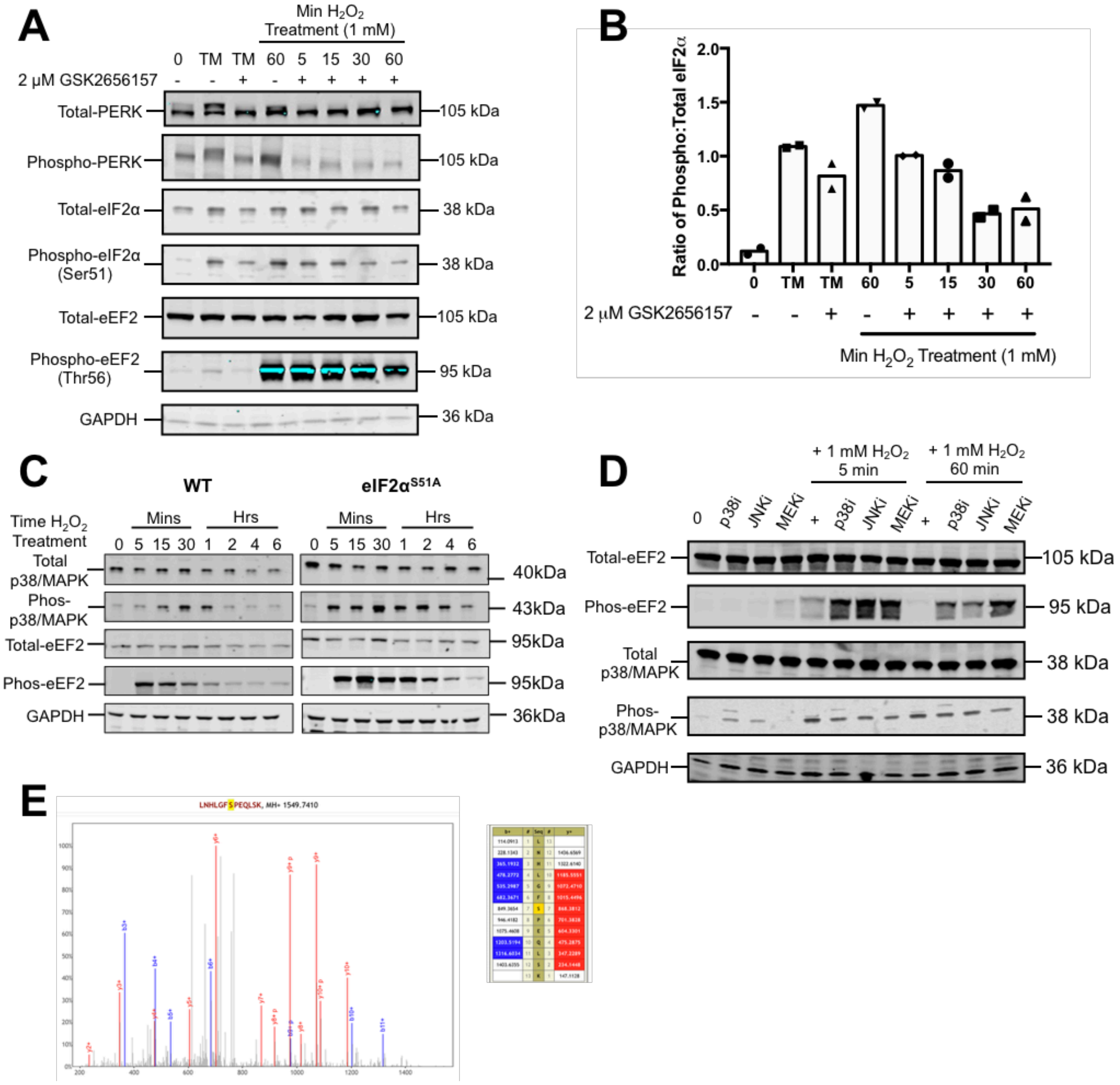


**Figure S6. Regulation of phosphorylation by different kinases in response to $H_2O_2$, Related to Figure 7.**

(**A**) Validation of PERK as the kinase that phosphorylates eIF2α under oxidative stress. WT MEFs were pre-treated for 1 hr with the PERK inhibitor GSK2656157 (2 μM) and as indicated and subsequently treated with 1 mM $H_2O_2$ for the indicated times. Western blot analysis was then performed.

(**B**) The graph represents quantification of Western blots from two separate experiments, including Figure S4A, using Licor imaging software. The individual symbols represent individual data points.

(**C**) The effects of p38/MAPK phosphorylation and inhibition on eEF2K under oxidative stress. Western blot analysis was done on WT and eIF2α$^{S51A}$ MEFs treated with 1 mM $H_2O_2$ for the time points indicated.

(**D**) WT cells were pretreated with different MAPK inhibitors (p38i: SB203580, 20 μM, JNKi: SP600125, 40 μM, MEKi: U0126, 50 μM) for 1 hr followed by treatment with 1 mM $H_2O_2$ for the indicated times. Western blot analysis was performed.

(**E**) Mass spectrometric evidence for $H_2O_2$-triggered phosphorylation of serine 436 of *S. pombe* Cmk2p. Wildtype cells were exposed to $H_2O_2$ for 15 min, followed by preparation of cell lysate for phosphoproteomics (Singec et al., 2016). The mass spectrum shows evidence of phosphorylation of serine 436.

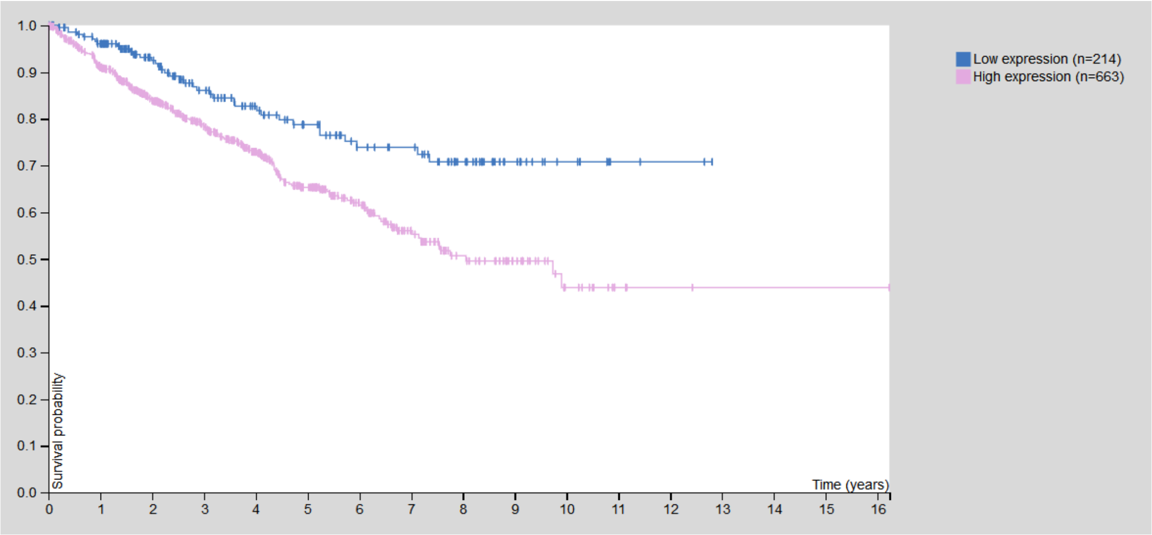


**Figure S7. High levels of EEF2K mRNA correlate with decreased survival in renal cancer, Related to Discussion.**

Kaplan-Meier plots summarizing the results from analysis of correlation between mRNA expression level and patient survival. RNA-seq data is reported as median FPKM (number Fragments Per Kilobase of exon per Million reads), generated by The Cancer Genome Atlas (TCGA). Patients were divided based on level of expression into one of the two groups "low" (under cut off) or "high" (over cut off). FPKM cut off = 3.51. P = 0.00027. Data obtained through www.proteinatlas.com.
